# Impact of Long-Term Mercury Contamination on the Rhizosphere Microbiota of *Lotus tenuis*: A Pathway to Resilience via Interkingdom Facilitation

**DOI:** 10.1101/2025.04.07.647525

**Authors:** Emanuela D. Tiodar, Cristina L. Văcar, Mihai C. Grimm, Iolanda V. Ganea, Zoltan R. Balázs, Anastasia M. Abrudan, Csaba Timár, Ioan Tanțău, Manuela Banciu, Roey Angel, Dorina Podar

**Affiliations:** Doctoral School of Integrative Biology, Babeș-Bolyai University, Cluj-Napoca, Romania; Centre for Systems Biology, Biodiversity and Bioresources (3B), Babeș-Bolyai University, Cluj-Napoca, Romania; National Institute for Research & Development of Isotopic & Molecular Technologies Cluj-Napoca; Department of Molecular Biology and Biotechnology, Babeș-Bolyai University, Cluj-Napoca, Romania; Babeș-Bolyai University, Cluj Napoca, Geology, Cluj Napoca, Romania; Institute of Soil Biology, Biology Centre CAS, České Budějovice, Czech Republic

**Keywords:** tolerance, phytoremediation, *Pseudomonas*, *Lotus tenuis*, *mercuric reductase* (MerA)

## Abstract

The rhizosphere microbial communities of *Lotus tenuis* in Hg-contaminated soils demonstrate remarkable resilience, maintaining stable bacterial and fungal diversity across a broad contamination gradient (40–1964 mg Hg kg⁻¹ soil). Despite significant shifts in community structure compared to control communities from uncontaminated rhizosphere soil, alpha diversity remained largely unaffected, likely due to widespread *merA*-mediated bacterial detoxification. These findings align with the stress gradient hypothesis, indicating that facilitative microbial interactions drive adaptation under Hg stress rather than competition-driven diversity loss. The rhizosphere was enriched with *Mesorhizobium* sp., supporting nitrogen fixation, *Pseudomonas* sp., a promising Hg-resistant bacterium, and various fungal partners enhancing plant tolerance to metal stress and drought. Key taxa included: *Streptomyces* sp. (a biocontrol agent), *Allorhizobium-Neorhizobium-Pararhizobium-Rhizobium* sp., *Shinella* sp. (rhizobial plant-growth promoters), *Nocardioides* and *Skermanella* potential metal- or aridity-resistant bacteria, *Darksidea* sp., *Acrocalymma paeoniae (*septate fungal endophytes that may promote plant stress tolerance), *Mortierella alpina, Chaetosphaeronema (*plant growth-promoting fungi)*, Humicola* sp., *Vishniacozyma* sp. (potential pathogen suppressors)*. Septoglomus*, *Dioszegia*, and *Articulospora* emerged as potential candidates for microorganism-assisted phytoremediation. This study provides a field-based, long-term perspective on microbial adaptation to Hg stress, highlighting plant-driven recruitment of beneficial microbiota as a key mechanism for ecosystem resilience and soil recovery.

## Introduction

Soil is the foundation of life on Earth, housing 59% of Earth’s species (Anthony et al., 2023) and supporting 95% of global food production (FAO, 2022), nourishing with healthy food and filtered water all of the aboveground biodiversity including the human population. Furthermore, soil mitigates the rapid advances of climate change through its carbon and water retaining properties - storing three times more carbon than the atmosphere (European Comission, 2011). Alarmingly, one-third of the world’s soil is already degraded (FAO & ITPS, 2015), and its degradation continues at a rate of 1 million square kilometers annually, surpassing the size of Antarctica (Tomalka et al., 2024). Soil degradation threatens global food security and ecosystem stability, and it leads to a costly loss of ecosystem services estimated between USD 6.3 - 10.6 trillion per year (Tomalka et al., 2024). In the European Union (EU) alone, more than 60% of soil-based ecosystems are unhealthy (European Comission, 2021) and it costs the EU 50 billion EUR per year (European Commission, 2020). Degradation causes are extensive and ubiquitous, from wind and water erosion to floods and landslides, loss of soil organic matter, salinization, contamination, compaction, sealing and loss of soil biodiversity (European Commission, 2016). Seeing how 70% of the world’s soil is in poor unhealthy conditions, restoring degraded soils has never been more urgent (European Commission, 2023).

Soil minerals are essential for plant growth as macro- and micronutrients (N, P, K, Ca, Mg, S, and Fe, Mn, Zn, Cu, Mo, respectively) (Marschner, 2012). Microorganisms, as well as plants, require minerals as cofactors for a wide variety of enzymes (Hemkemeyer et al., 2021). However, a disproportionate accumulation of trace metals in soils can lead to serious contamination issues and deleterious biodiversity effects. Over 30% of the European agricultural land area is polluted with heavy metals, the group of metals that typically have a density of more than 5 g cm^-3^ (Prăvălie et al., 2024). Some trace metals are not part of the cellular metal homeostasis network and do not serve a biological function (e.g. Cd, Hg, Pb). Moreover, under elevated concentrations, trace metals are poisonous and can induce toxicity symptoms to plant and microbiota (Abdu et al., 2017).

Mercury and Hg compounds are listed among the top 20 priority hazardous substances for EU water bodies (Water Frame Directive, 2008) and Hg ranks third on the substance priority risk for human health (ATSDR, 2022) due to its persistence in the environment and biomagnification within the food chain. Moreover, Hg pollution is a significant global threat to soils, with anthropogenic releases contributing approximately 13% of the total Hg in organic soil layers over the natural background of Hg in land (AMAP/UN, 2019).

Mercury is toxic to plants, symptoms of toxicity including: chlorophyll degradation, protein metabolism disruptions, aberrant reactive oxygen species (ROS) generation, and in legumes, reduction in the number or size of nodules (Natasha et al., 2020; Quiñones et al., 2013; Safari et al., 2019). Thus, Hg-contaminated sites tend to lack vegetation, while scarce pioneer species slowly establish (Muryani et al., 2023; Tiodar et al., 2024). Remarkably, plants have developed Hg tolerance strategies stemming from the mechanisms of the metal homeostasis network (Tiodar et al., 2021). Some plants are able to reduce Hg uptake by binding the mercuric ion through root exudates, thiol groups found in the root cell wall, or to the hemicellulose in the leaf cell walls for atmospheric Hg deposition (Carrasco-Gil et al., 2013; Montiel-Rozas et al., 2016; Sun et al., 2021), whereas others sequester Hg in vacuoles after intracellular chelation by macromolecules such as phytochelatines (Park et al., 2012; L. Sun et al., 2018). To date, no plant species has been identified as truly Hg-tolerant able to accumulate high amounts of Hg in shoots via (soil-to-)root translocation. However, native pioneer species capable of thriving in Hg-contaminated environments, along with their associated rhizosphere microbiota, represent a valuable reservoir of bioresources. These plant-microbe associations hold significant potential for the development of nature-based solutions for Hg remediation, either as a primary strategy for large-scale contaminated sites or as a complementary approach following mechanical interventions. Among the native plant species growing in Hg-contaminated soils, legumes are of interest due to their root symbiosis with bacterial species able to fix nitrogen, thereby naturally fertilizing soil with nitrogen for the benefit of future plants.

Short-term Hg exposure reduces microbial diversity and significantly alters community structure (Frey & Rieder, 2013; Frossard et al., 2017; Zheng et al., 2022), though fungal communities are generally more resistant than bacteria (Durand et al., 2020; Frossard et al., 2017). However, under long-term exposure, microbial communities gradually adapt, recovering diversity even under high Hg concentrations (Frossard et al., 2018; Liu et al., 2014). Over time, microbial composition in contaminated soils is shaped more by other edaphic factors (e.g., pH, organic matter) than by total Hg soil concentration (Liu et al., 2014; Zheng et al., 2022). Therefore, we assume that microbial responses to Hg stress follows two stages: an initial decline in diversity due to the loss of Hg-sensitive taxa, and a subsequent recovery as resistant taxa proliferate, mitigating stress to a level acceptable for the re-establishment of sensitive species through facilitative interactions (Durand et al., 2020). However, decisive field studies tracking these composition changes over time are lacking. Understanding how microbial communities restructure under chronic Hg exposure in natural settings remains a key research gap. Nevertheless, community resilience may emerge through the acquisition of Hg resistance traits and mechanisms.

The common pathways employed in Hg resistance by both bacteria and fungi are cell-wall binding to limit Hg uptake (Kinoshita et al., 2013; Puglisi et al., 2012) and intracellular chelation by peptides and proteins rich in sulfhydryl (-SH) groups to limit Hg bioavailability (Y. Guo et al., 2025; Lima et al., 2006; Manceau et al., 2021). Most importantly, bacteria can employ Hg detoxification through biovolatilization, mechanism, encoded by the *mer* operon. The key enzyme of the *mer* system is the mercuric reductase encoded for by the *merA* gene, that converts Hg^2+^ to the inert, volatile form of Hg^0^. The *merA* gene is ubiquitously found in both the chromosome of diverse bacterial phyla (e.g. Pseudomonadota (Proteobacteria), Bacillota (Firmicutes), Actinomycetota, Aquificota, Bacteroidota, Chloroflexota, Deinococcota, Mycoplasmatota, Nitrospirota, and Verrucomicrobiota) and on plasmids that can be more easily shared between phylogenetically distinct taxa/bacteria via lateral gene transfer (Boyd & Barkay, 2012; Christakis et al., 2021). For fungi, intracellular chelation of Hg ions could be performed by glutathione, metallothioneins, phytochelatins (Bolchi et al., 2011; Lorenzo-Gutiérrez et al., 2019; Raspanti et al., 2009), and even selenium rich proteins (Kavčič et al., 2019). Moreover, fungi can also rely on a tolerance mechanism of compartmentalization by transferring the glutathione-chelated complexes into the vacuoles potentially through transporters such as Ycf1 (Gueldry et al., 2003). Nonetheless, Hg volatilization has also been observed for fungi, albeit the molecular determinants have not been fully characterized, the *merA*-mediated system of Hg reduction seems to be widely distributed (Chang et al., 2021; Pietro-Souza et al., 2020; Urík et al., 2014; Wu et al., 2022).

Finding Hg resistant bacterial and fungal species is the key to designing nature-based solutions for restoring contaminated lands that combine microbiota assistance to plant metal stabilization. Mercury resistant isolates of *Rhizobium leguminosarum*, *Bradyrhizobium canariense*, and *Ensifer meliloti* have been previously isolated from legume root nodules in the Hg-contaminated mining area of Almadén, Spain (Ruiz-Díez et al., 2012). These isolates exhibited both *in vitro* Hg tolerance and nodule-forming capacity. Under controlled conditions, an Hg-tolerant *B. canariense* strain conferred Hg tolerance to a sensitive *Lupinus albus* cultivar grown in a substrate containing up to 102 mg Hg kg⁻¹, without reducing shoot biomass or chlorophyll content (Quiñones et al., 2013). The plants accumulated high Hg levels in roots and nodules, promoting root Hg stabilization. Notably, only Hg-tolerant bacterial strains conferred this benefit, as plants inoculated with Hg-sensitive strains did not exhibit tolerance.

The identification of Hg-tolerant bacterial strains is crucial for advancing microorganism-assisted phytoremediation. This strategy leverages plant-microbe interactions to enhance plant growth, biomass yield, and contaminant stabilization, while also providing key ecosystem services such as pollutant degradation, nutrient cycling, and plant growth promotion (Saccá et al., 2017). Notably, both tolerant and sensitive native plants can be enhanced for Hg phytoremediation through inoculation with single strains or microbial consortia, whether native or introduced.

In this study, we aimed to investigate the long-term (>30 years) effects of a Hg contamination gradient on the rhizosphere soil microbiota associated with *Lotus tenuis*. Previously, *L. tenuis* was identified as an abundant legume pioneer colonizer of a highly Hg-contaminated brownfield, being able to immobilize on average 4479 mg Hg kg^-1^ root in soils with an average concentration of 1308 mg Hg kg^-1^ (Tiodar et al., 2024). Herein, we focused on the bacterial and fungal Hg-tolerant rhizosphere communities, assessing shifts in community structure response to a Hg rhizosphere soil concentration gradient and, in comparison to uncontaminated control soils. We hypothesized that facilitative interactions within microbiota communities would have stabilized the microbial diversity and functionality under the Hg gradient, according to the principles of the stress gradient hypothesis (Bertness & Callaway, 1994). Therefore, by characterizing the microbial communities, we aimed to (1) assess the community resilience under the Hg gradient and detect potential resistance mechanisms employed at the community level, (2) identify (differentially abundant) taxa associated with Hg contamination within rhizosphere soils, and (3) identify taxa specifically associated to plants with higher Hg accumulation potential.

## Materials and Methods

### 1. Site description

The two sites where sampling occurred were located approximately 4 km apart, in the area of the Turda City, Cluj County, Northwest Romania (Fig. 1). The first site of approx. 6000 m^2^ (N = 46.558389°, E = 23.779794°), on the grounds of the former Turda Chemical Plant consists of an industrially contaminated brownfield where a chlor-alkali production facility was active for more than 30 years (from here on referred to as Hg site). In the “chlor-alkali” process, chlorine gas (Cl_2_) was generated *via* sodium chloride (NaCl) electrolysis. This electrochemical reaction was catalysed by Hg cathods (Maghear, 2013). Since 1958, activity at the site ceased after the chlor-alkali production unit was decommissioned. Subsequently, the former unit was abandoned and the building was demolished without taking proper contamination containment measures and without soil remediation and restoration procedures. Today, the main contaminant of the soil is Hg, and the contamination levels at the site are highly heterogeneous. Recently, the median concentration value at the site was estimated at 962 mg Hg kg^-1^ topsoil (Tiodar et al., 2024), which exceeds the national industrial soil legal limit by almost 100-fold (Ministerial Order, 1997). Moreover, Pb, Cu, Zn, and Mn are also soil co-contaminants at different risk degrees. The area is dominated by cement debris, and characterized by high concentrations of toxic metals in the surface layers of the soil. Thus, this brownfield is an open source of metal contamination for the urban population in near vicinity through wind (Esbrí et al., 2018) and land erosion, with an additional risk of metals leaching into the nearby Arieș River. Nevertheless, plant pioneers are colonizing the area, even the Hg hotspots, but the plant coverage is limited, patchy, and the damaged background of the area hinders floral succession (Tiodar et al., 2024). The second site (N = 46.587155°, E = 23.792483°), a typical densely vegetated temperate meadow, mostly represented by grass species, was selected for this study as a reference, since there is no known metal contamination (referred to as Control site from here on).

**Fig. 1.**
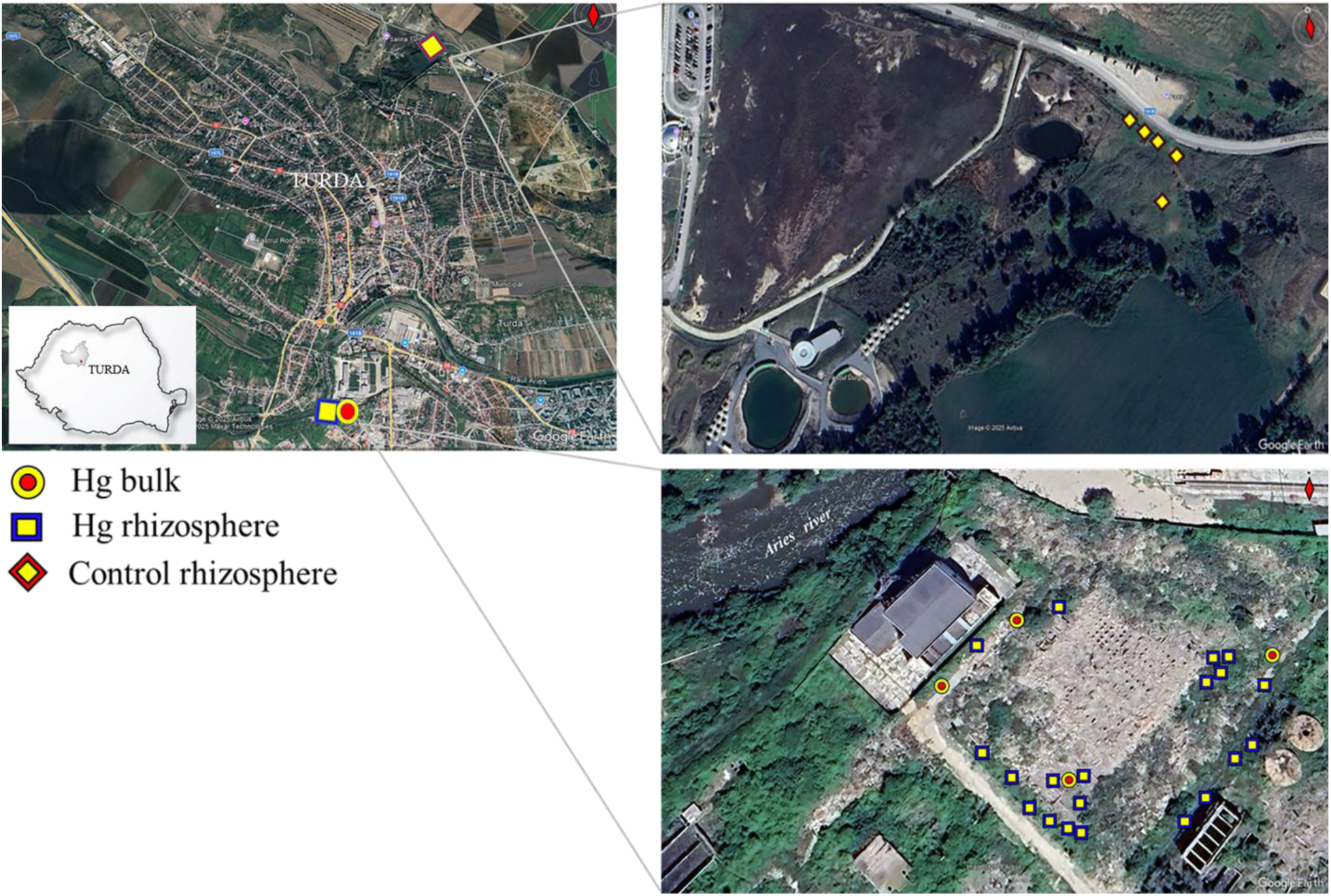
Maps (GoogleMap, 2025) of the two study sites with the sampling points. The industrial brownfield, Hg site (bottom) with the 20 rhizosphere (blue squares with a yellow center) and 4 bulk (b) soil sampling points (yellow circles with a red center), and the Control greenfield site (top) with 5 rhizosphere (c) soil sampling points (red diamond shapes with yellow center) superimposed on the map.

### 2. Material collection

On May 15, 2023, both sites were sampled for plant and soil samples. *Lotus tenuis* Waldst. & Kit. ex Willd (Narrowleaf trefoil; a common N-fixing pioneer leguminous perennial) individuals were identified as per voucher numbers 674507 (Hg site) and 674508 (Control site) deposited at the Herbarium Musei Universitatis Napocensis, photographed, and collected from both sites (twenty individual samples at the Hg site and five at the Control site). Plants were separated into roots and shoots, shoot fresh weight (FW) was recorded, and the material was placed in individual plastic bags and transferred on ice to the laboratory. Rhizosphere soil samples collected together with the roots, were placed in plastic bags on ice, and were only manually separated from the roots in the laboratory after several hours. Rhizosphere soil was considered in this study as the soil closely adhering to the roots that did not come off after a few seconds of manually shaking the roots in the field. The rhizosphere soil fraction included also symbiosis nodules displaced from the roots during collection. These rhizosphere soil samples were sieved of leftover fine roots and mixed prior to being flash frozen in liquid N_2_ and stored at -80 °C until DNA extraction was performed. From underneath the root system of each plant individual, bulk soil (referred to as plant bulk soil) was also collected into plastic bags, placed on ice, and stored at 4 °C until further physico-chemical processing. At the Hg site, four bulk soil samples from unvegetated sampling points were also collected, and in the laboratory, the soil was sieved through a 2 mm mesh and a subsample was treated in the same way as the rhizosphere soil samples to be used for DNA extraction. In the laboratory, roots and shoots were thoroughly washed under running cold tap water, then rinsed with distilled water, to ensure that leftover soil particles or dust were removed from the material surface. After blotting, plant tissue was preserved in plastic bags at -20 °C to avoid Hg losses until elemental determination.

### 3. Metal concentration measurements

Plant bulk and unvegetated bulk soil samples were sieved through a 2 mm mesh. Fresh weight was recorded, then samples were left to dry at 30 °C for 2 d to avoid Hg evaporation. After overnight room temperature acclimatization, dry weight (DW) was measured. Dried samples were ground to fine powder with agate mortar and pestle. Powdered soil samples (0.25 g) were added to a Teflon vessel and mixed with aqua regia (HCl, ≤10^-4^% Hg, HNO_3_, ≤5•10^-4^% Hg, Merck, Darmstadt, Germany) prior to microwave digestion (Multiwave 5000, Anton Paar, Graz, Austria) at 180 °C for 30 min, following a leaching protocol based on EPA 3051A (US EPA, 2007). Shoots and roots samples were dried at 30 °C for 14 days, and the material was ground in porcelain mortar and pestle. The ground tissue (0.4 g) was digested in a HNO_3_:HCl mixture (1:2, v/v) for 20 min at 180 °C in the microwave digestor. An inductively coupled plasma-optical emission spectrometer PlasmaQuant 9100 Elite High Resolution ICP-OES (Analytik Jena, Jena, Germany) equipped with an Echelle double monochromator, a concentric borosilicate nebulizer, a borosilicate glass cyclonic spray chamber and a V Shuttle Torch with 2 mm quartz injector was used for routine analysis of chemical elements in the digested samples. The spectrometer was coupled to a Cetac ASX 560 Autosampler (Teledyne CETAC Technologies, Omaha, NE, USA) with 240 positions. The axial and radial configurations were employed to identify the elements at the following spectral lines: Mn II 257.610 nm, Zn II 206.200 nm, Cu I 324.754 nm, Fe II 259.940 nm, Pb II 220.353 nm, Mg I 285.213 nm, Ca II 315.887 nm, and Hg II 194.159 nm. Hg measurements were conducted after prior complexation of all samples with a 1 mg/L solution of 1000 mg/L Gold standard TraceCERT® (Sigma Aldrich, St. Louis, MO, USA). All measurements were performed in triplicate, 2 QC standards, Standard Reference Material 1547® (SRM 1547) Peach Leaves (NIST, Gaithersburg, MD, USA), SRM 2710a Montana II soil (NIST, Gaithersburg, MD, USA), and blanks were prepared for the analysis. The standard’s measured element concentrations, recovery and correction factors calculated for each metal species determined are shown in Supplementary Table 1. All metal concentration values are given on a dry weight (DW) basis. Based on the Hg concentrations in the samples, a virtual gradient was used as a guide to further study the microbial Hg resistance.

### 4. Soil analyses

Soil subsamples were used for pH, moisture, total organic carbon (TOC), and carbonates content analyses. The pH was measured in a suspension prepared by adding 2.4 g soil in a 6 mL 0.01M CaCl_2_ solution and vigorous mixing for a few seconds. Moisture content was estimated after drying the samples at 105 °C for 12 hours. For estimating TOC and carbonate content, a loss on ignition protocol was followed, with repeated mass weighting after subsequent oven drying at 550, and 950 °C, respectively.

### 5. Soil DNA Extraction and 16S rRNA and ITS gene sequencing

Total soil DNA was extracted in triplicates from 0.25 g rhizosphere and unvegetated bulk soil samples using the Quick-DNA**™** Fecal/Soil Microbe Miniprep Kit (ZymoResearch, #D6010). Mechanical lysis was performed by vigorous shaking (35 Hz) for 45 minutes on a Vortex Genie^®^. All other steps were conducted following the instructions provided by the kit manufacturer. DNA concentration and quality was checked by Nanodrop (ThermoFisher Scientific), Qubit (Invitrogen), and gel electrophoresis after which the samples were shipped to Macrogen, Inc. Europe. Further 16S rRNA V3-V4 and ITS fITS7-ITS4 library preparations and quality checks were performed prior to sequencing on an Illumina MiSeq platform with 300 bp paired-end reads. The primer pairs used for generating the 16S and ITS libraries were, respectively, the universal 341F (CCTACGGGNGGCWGCAG) - 805R (GACTACHVGGGTATCTAATCC) and fITS7 (GTGARTCATCGAATCTTTG) (Ihrmark et al., 2012) - universal ITS4 (TCCTCCGCTTATTGATATGC). The raw reads sequence data have been deposited in the European Nucleotide Archive (ENA) at EMBL-EBI under project accession numbers PRJEB88147.

### 6. Endpoint PCR targeting the bacterial *merA* gene

The presence/absence of the *merA* gene was checked using endpoint PCR in the same total soil DNA extracts as for the metabarcoding procedure. The *merA* gene was selected as a marker for Hg resistance to determine the acquisition of functional traits by the bacterial communities across the rhizosphere soil Hg gradient. Degenerate primer sets (merA7sn_F and merA5n_R: as specified in Liu et al., 2012) were used for a broader targeting strategy as *merA* nucleotide sequence can vary between bacterial phyla. The reaction conditions were previously reported in Tiodar et al., 2024.

### 7. Sequence data processing

Demultiplexed 16S rRNA gene sequencing FASTQ raw data files were first put through the cutadapt function to remove the primer sequences, then filtered and trimmed based on the error rate algorithm computed by the DADA2 pipeline (Callahan et al., 2016). After sequence inferring, the two pair-end reads were merged into individual amplicon sequence variants (ASV). The merged sequences were of varied lengths ranging from 292 to 464, with two peaks at 402 and 427. Next, denoising and chimera removal was performed, and the true biological sequences were assigned taxonomy based on the SILVA database (ver. 138, silva_nr99_v138_wSpecies_train_set.fa.gz) prior to checking for contamination using decontam (V1.24; Davis et al., 2018).

The demultiplexed paired-end ITS reads were loaded into the PIPITS pipeline (ver. 3.0) as FASTQ files (Gweon et al., 2015), where the read-pairs were merged, quality filtered and dereplicated. Next, the fungal ITS2 subregion from each merged read was extracted, and clustered into operational taxonomic units (OTU) (at 97% identity) prior to subjecting the resulting representative sequence to chimera removal. However, to assign taxonomy against the UNITE fungal ITS reference database (UNITE_v10_21.04.2024; Abarenkov et al., 2024) the PIPITS output FASTA sequence files were manually searched using BLASTn (using a custom made bash script) against the UNITE database. The OTU sequences were checked for artifacts and manually curated based on the following guidelines: the sequences with less than 80% coverage were removed, all hits with e-value larger than 1e-20 were removed, and taxonomic assignment was performed up to genus level for queries with %identity ≥90, up to family for %identity ≥85, and up to order for %identity ≥75.

The resulting 16S and ITS abundance datasets were decontaminated based on frequency of contaminants (Davies et al., 2018), and library sizes were further reduced according to a sequence prevalence threshold of 5%. Library sizes dimensions for the two datasets can be checked in Supplementary Table 2 and 3.

Next, the effect of library sizes on community composition was checked with a PERMANOVA at 9999 permutations using the adonis2() function (*vegan* package; Oksanen et al., 2017). Based on the non-significant *p*-value (>0.5) and low R^2^ we proceeded further without library normalization to calculate alpha- and beta-diversity metrics (Supplementary Table 4). All data was visualized and analyzed in R ver. 4.3.1 (R Core Team, 2024).

### 8. Statistical analyses

For the environmental variables (moisture, TOC, metal concentrations), testing for differences in the means of the three groups defined by condition (rhizosphere vs. bulk) and treatment (Hg-contaminated site vs. Control site) was performed with one-way analysis of variance (ANOVA), followed by a posteriori comparison using Tukey’s test for unequal sample sizes (Sokal & Rohlf, 2012). To fit the ANOVA assumption of normal distribution, all environmental variable data were logarithmically transformed, except for the pH values. Thus, variances between groups were homogenized (Levene’s test for medians) and normal distribution was visualized through histograms, the Shapiro-Wilk test, and normal probability plots. For the plant variables (DW and metal concentrations), independent two-sample t-tests for equal or unequal variance were used to detect differences in the means of the two groups separated by site (Hg-contaminated, n=20 vs. Control, n=5).

The microbial communities were compared using alpha- and beta-diversity metrics based on the ASV and OTU counts. We used the observed number of species (S.obs), Chao1 and Abundance-based Coverage Estimator (ACE) indices to determine within sample differences in community richness, and the Shannon, Inverse Simpson and Berger Parker (BP) indices for diversity estimations (R *vegan* package, V2.6-6.1; Oksanen et al., 2017). General linear models (GLM) were used to test for differences in the estimators among groups, after assessing the heteroscedasticity between groups (Levene’s test for medians) (ST5). Pearson correlation coefficients were calculated to determine potential linear relationships between alpha diversity estimates and the Hg gradient. To visualize the dissimilarity in the structure of the microbial communities, the multidimensionality of the data was reduced based on the metric- (Principle Coordinate Analysis (PCoA)) and non-metric (NMDS) Multidimensional Scaling algorithms performed on Bray-Curtis distance matrices. Observed between groups differences were tested for significance with a (simple and pair-wise, according to the case) PERMANOVA at 9999 permutations, using the adonis function from the *vegan* R package. Based on the significant PERMANOVA results, constrained PCoAs were generated using the significant factors as constraining variables. Moreover, the Bray-Curtis distance matrices were used as the response variables characterizing the microbial community structure in distance-based Redundancy Analysis models (db-RDA) computed to determine the main environmental drivers of the community structures. A forward-backward selection of the variables was performed to select a significant parsimonious model using the ordiR2step function (*vegan*). The environmental variables were added onto the model and further tested for significance using the envit() function (*vegan*). The parsimonious models and their axes were tested for significance using ANOVA with 999 permutations. The Analysis of Compositions of Microbiomes with Bias Correction 2 (ANCOM-BC2) method (H. Lin & Peddada, 2020) was used to identify differentially abundant ASVs/OTUs among the three soil microbiomes categories (Hg bulk, Control rhizosphere, Hg rhizosphere). The analysis was performed using the *ancombc2()* function with soil type (Hg rhizosphere vs. Hg bulk vs. Control rhizosphere) as the sole factor and 100 bootstrap replicates, on raw microbial community data with a prevalence cut of 10%. The *p*-values were corrected according to the Benjamini-Hochberg (BH) adjustment and taxa were considered significantly different if *p* < 0.05.

The fungal OTUs with >97% similarity to the reference sequences of the UNITE database were further classified into ecologically relevant guilds/ trophic modes according to the FungalTraits database (Põlme et al., 2021). Only 13.4% of OTUs were assigned a fungal guild/ trophic mode. The relative abundance of the identified guilds and trophic modes was calculated per each community to determine any shifts in the functional pattern due to the soil Hg gradient. Statistical outputs, full ANOVA tables, and guild abundance comparisons are included in the Supplementary Information (Zenodo: https://doi.org/10.5281/zenodo.15165625).

## Results

### Plant fitness and Hg content

The two *Lotus tenuis* populations, from the Hg-contaminated (Hg site) and the Control site, were in their vegetative growth, pre-flowering stage, with no visible signs of toxicity at the sampling time. Fewer individuals were present within the population at the Control site in comparison to the Hg site. At Hg site, the plant population was distributed mainly towards the margins of the investigated area, where colonization was not hindered by concrete debris, as seen in Fig. 1. The plant dry biomass (DW) of individuals at the Hg site was generally higher than of the individuals at the Control site (Fig. S1a and b). With respect to Hg accumulation in plant tissues, the concentration of Hg in root and shoots of individuals from the Control site was negligible (Fig. S1), whereas, the concentration of Hg in roots and shoots of *L. tenuis* individuals at Hg-site, was dependent on, and positively correlated with metal concentration in the soil (Fig. S2).

### Edaphic Parameters

Soil moisture was significantly lower at the Hg site than at the Control site (Fig. 2a). No significant differences were determined in soil moisture between the rhizosphere and the bulk soil, at the Hg site (Fig. 2b). Soil pH was slightly alkaline ranging between 7-7.7, without any significant differences between the two sites or between bulk and rhizosphere. The median total organic carbon content in the rhizosphere soil at Control site was statistically significantly 1.5 and 2 times higher than in the rhizosphere and bulk soil, respectively, at the Hg-site (Fig. 2c). Carbonates content was similar in both soil types (bulk vs. rhizosphere) and sites (Control vs. Hg-site) (2d).

**Fig. 2.**
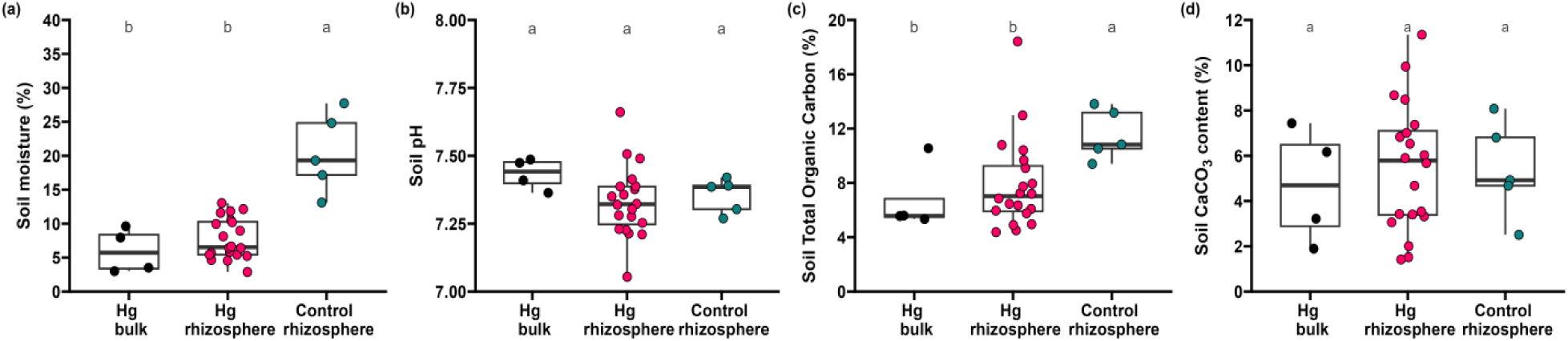
Environmental soil conditions: moisture (a), pH (b), total organic carbon (c), and CaCO_3_ content (d) at the Control and Hg-contaminated sites, in Turda, Romania. The boxplots show the median (horizontal line inside box), the interquartile range (IQR), and the individual data points (circles). All environmental variables, except soil pH, were log-transformed prior to an ANOVA test. Different letters denote significant differences between means as determined by *a posteriori* Tukey’s test. n = 4 – 20.

The concentrations of plant macroelements in soils did not vary significantly between the two sites (Fig. S2a), and neither between the bulk and *L. tenuis* rhizosphere soil. The same could be observed for the plant microelements (Fig. S2b). However, the concentrations of the toxic elements, Hg and Pb were significantly higher at Hg site compared to the Control site. The median Hg concentration in bulk and rhizosphere soil was 22 and 50 times higher, respectively at Hg compared to Control site(Fig. 3).

**Fig. 3.**
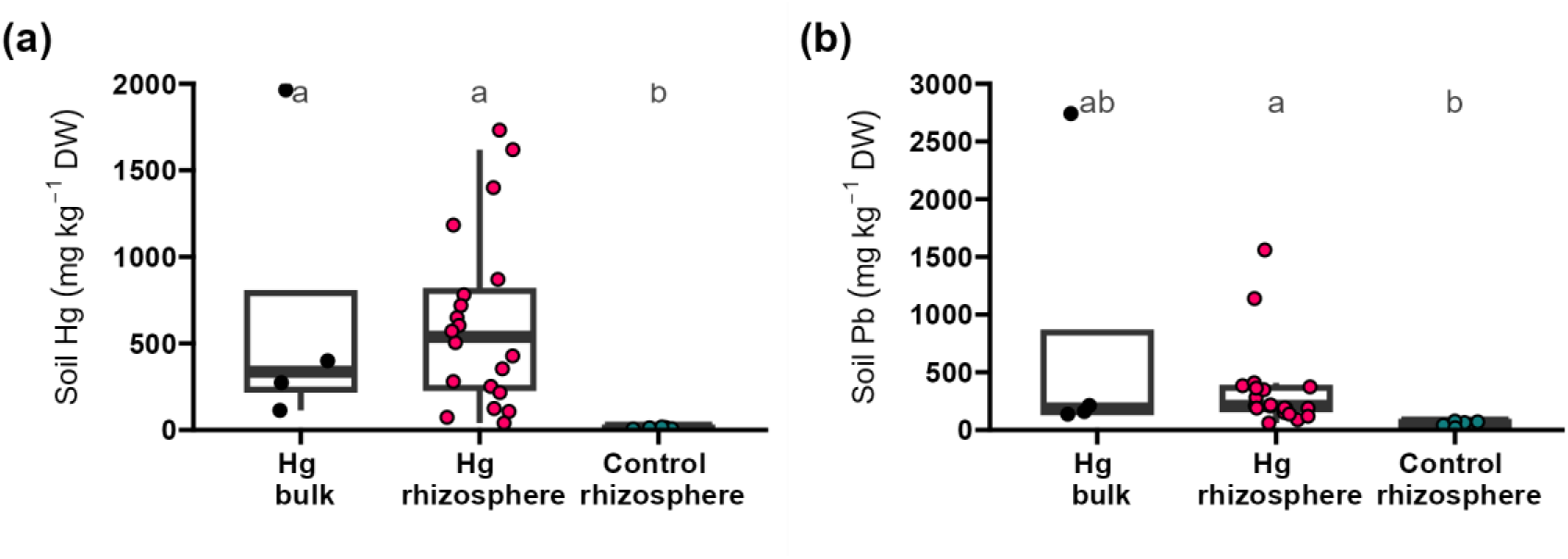
Concentrations of toxic elements in soil: mercury (a) and lead (b) at the Control and Hg-contaminated sites, in Turda, Romania. The boxplots show the median (horizontal line inside box), the interquartile range (IQR), and the individual data points (circles). Soil metal concentrations were log-transformed prior to an ANOVA test. Different letters denote significant differences between means as determined by *a posteriori* Tukey’s test. n = 4 – 20.

### 2. Microbiota community structure

#### Alpha diversity - Bacteria (and Archaea) and fungi

The alpha diversity of *L. tenuis* associated rhizosphere bacterial and fungal communities was examined in the absence of Hg, along a gradient of 40-1732 mg Hg kg^-1^ soil, and for Hg-contaminated bulk soil (113-1964 mg Hg kg^-1^ soil) (Fig. 4, and Fig. S3). Largely, alpha diversity indices of rhizosphere bacterial and fungal communities were not linearly dependent on the soil gradient of Hg concentration (Fig. S4). In particular, community richness was not significantly different for neither soil condition (bulk vs. rhizosphere) nor for site, for neither the bacterial, nor fungal communities (Fig. 4). Significant diversity differences were observed as per the Shannon Diversity index and the Inverse Simpson estimator. Bacterial bulk and rhizosphere communities at the Hg-contaminated site exhibited lower Inverse Simpson estimators compared to the Control rhizosphere communities, with the bulk communities having a 1.5-fold decrease compared to the Control (Fig. 4a). Moreover, the Hg rhizosphere fungal communities exhibited the lowest Inverse Simpson index, which had a significant 2-fold lower median value than the control and the bulk communities (Fig. 4b).

**Fig. 4.**
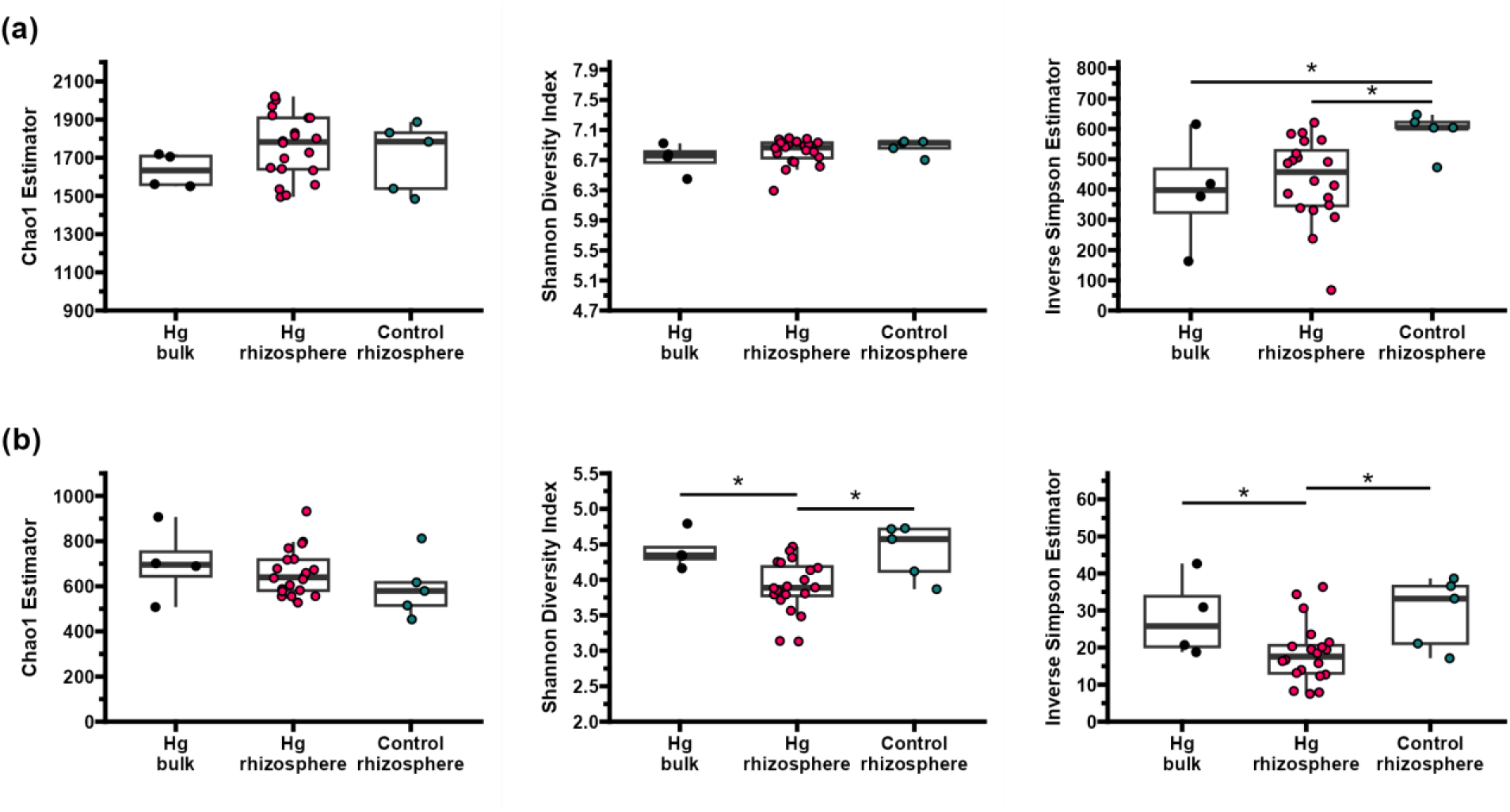
Alpha diversity of microbial rhizosphere and bulk communities in the absence of Hg (Control) and under Hg stress (Hg). Boxplots show the Chao index, Shannon index, and the Inverse Simpson estimator for bacteria (a) and fungi (b). The asterisk denotes significant differences (*p* < 0.05) between conditions according to a generalized linear model that was used to compare each alpha metric of a soil condition versus the other two conditions (*p* < 0.05).

#### Profiles of microbial phylum/order abundance between sample types

The first three most abundant bacterial phyla at both sites were Actinomycetota (Actinobacteriota), Pseudomonadota (Proteobacteria), and Acidobacteriota (Fig. 5). At the Hg site, the relative abundance proportions of Actinomycetota to Pseudomonadota were equal for rhizosphere soil. However, at the Control site, Actinomycetota had a 1.5-fold mean relative abundance increase compared to Pseudomonadota, in the rhizsophere soil. Among the two sites, Bacillota (Firmicutes) had a 3.6-fold mean relative abundance decrease at the Hg site compared to the Control site. In the rhizosphere soil at the Hg site, the relative abundance of Bacteroidota was approximately 2 times higher than in bulk soil or in rhizosphere soil at the Control site. The Myxococcota phylum had approximately twice as high mean relative abundance in rhizosphere soils than in bulk soil.

**Fig. 5.**
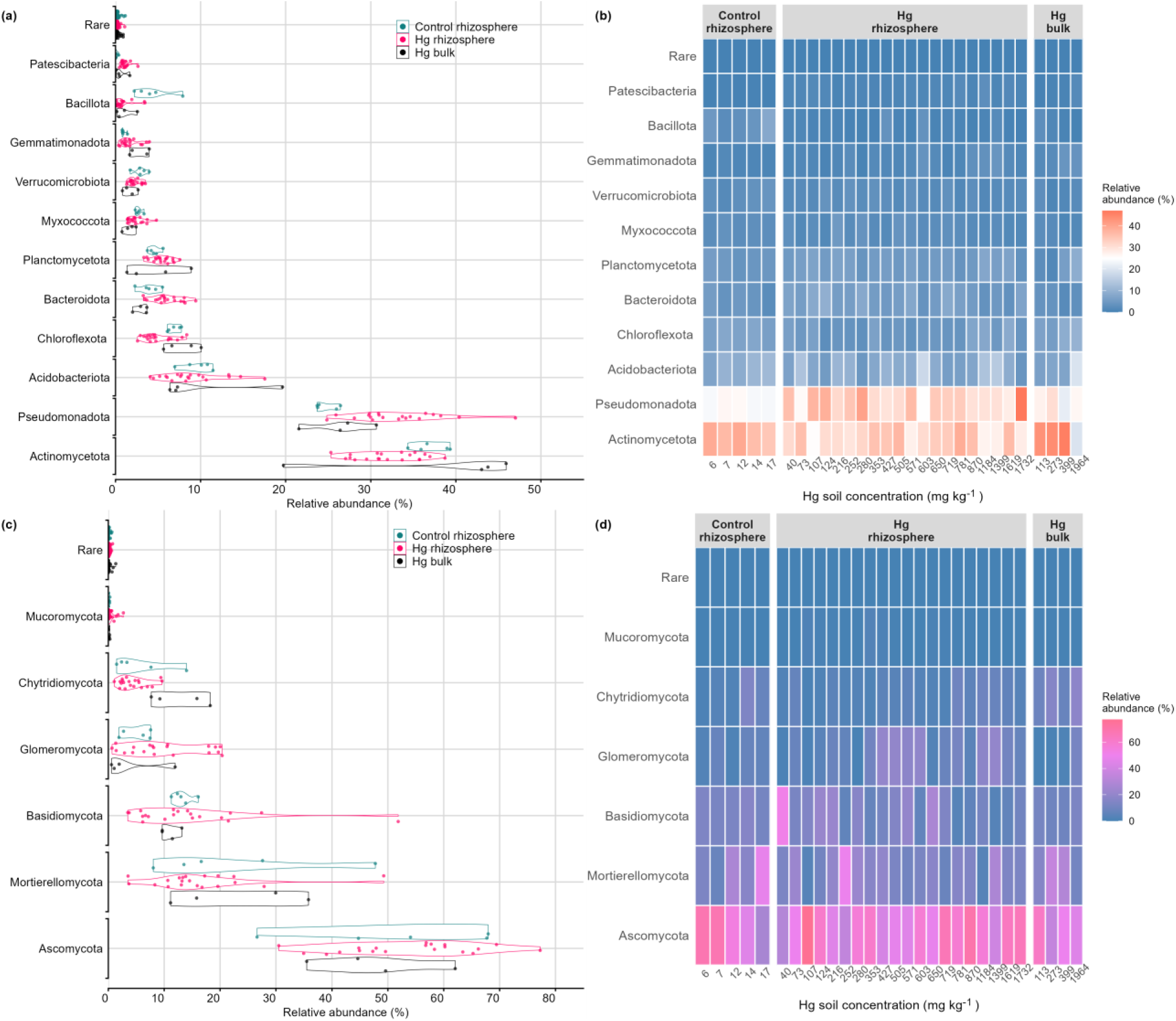
Violin and heatmap plots of the phylum relative abundance of the bacteria (a, b) and fungal (c, d) communities. Rare phyla were grouped together if the relative abundance was under 0.01% across all samples.

The four most abundant bacterial orders in the 16S dataset were Rhizobiales, Solirubrobacteriales, Propionibacteriales, and Burkholderiales, and the relative abundance of these orders did not differ greatly between the Hg rhizosphere, Hg bulk, and Control soil communities (Fig. S5). At the Hg site, the mean relative abundance of Rhizobiales was 1.6 times less in the bulk communities compared to the rhizosphere soil communities. The Hg communities (both rhizosphere and bulk) had a 6-, 3-, and 2-fold mean relative abundance decrease in Bacillales, Gaiellales, and Vicinamibacterales, respectively and a 5-fold increase in Frankiales compare to the mean relative abundance of the Control rhizosphere communities.

The most abundant fungal phyla were Ascomycota, Mortierellomycota, and Basidiomycota (Fig. 5). Overall, the three phyla comprised 88% in the Control rhizosphere soil, 85% in the Hg rhizosphere soil, and 82% in the Hg bulk soil of the total group mean relative abundance. Chytridiomycota had a mean relative abundance similar to Basidiomycota in the Hg bulk soil, where it was on average 3 times higher than in the rhizosphere soil samples. Glomeromycota was two times more abundant in rhizosphere than in bulk soil, and between the rhizosphere communities of the two sites, it had a two-fold increase in mean relative abundance at the Hg site compared to the Control site. Rozellomycota had a 2-fold mean relative abundance decrease respectively at the Hg site compared to the Control site.

The most abundant fungal Order in the Hg rhizosphere communities was Pleosporales, with a 3 times higher relative abundance than the Control site communities (Fig. S5). Moreover, Glomerales had a 3-fold, Pezizales, and Filobasidiales a 2-fold increase in relative abundance in the Hg rhizosphere soil than in Control or Hg bulk soil. Hg sensitive fungi belonged to the Hypocreales and Chaetothyriales orders, which had a 2- and 4-fold decrease respectively in relative abundance at the Hg site compared to the Control.

The ANCOM-BC2 algorithm was used to identify taxa associated to Hg resistance in the *L. tenuis* rhizosphere. Thus, 24 bacterial and 58 fungal taxa were found to be globally significant when comparing the Hg-contaminated rhizosphere communities to the Control and Hg bulk soil communities in a pairwise fashion at the ASV/OTU level (Fig. 6). Half of the globally significant bacterial taxa were differentially more abundant in the Hg-contaminated rhizosphere than in both the Control rhizosphere and Hg bulk communities. Among the more abundant bacterial genera in the L. tenuis rhizosphere at the Hg-contaminated site were: *Streptomyces* sp., *Steroidobacter* sp., *Skermanella* sp., *Shinella* sp., *Pseudonocardia* sp., *Nocardioides* sp., *Iamia* sp., *Cellulomonas* sp., *Allorhizobium-Neorhizobium-Pararhizobium-Rhizobium* sp., and *Acidibacter* sp. On the Hg-sensitive side, *Reyranella* sp., *Nordella* sp., and *Flavisolibacter* sp. were more abundant in the Control rhizosphere.

**Fig. 6.**
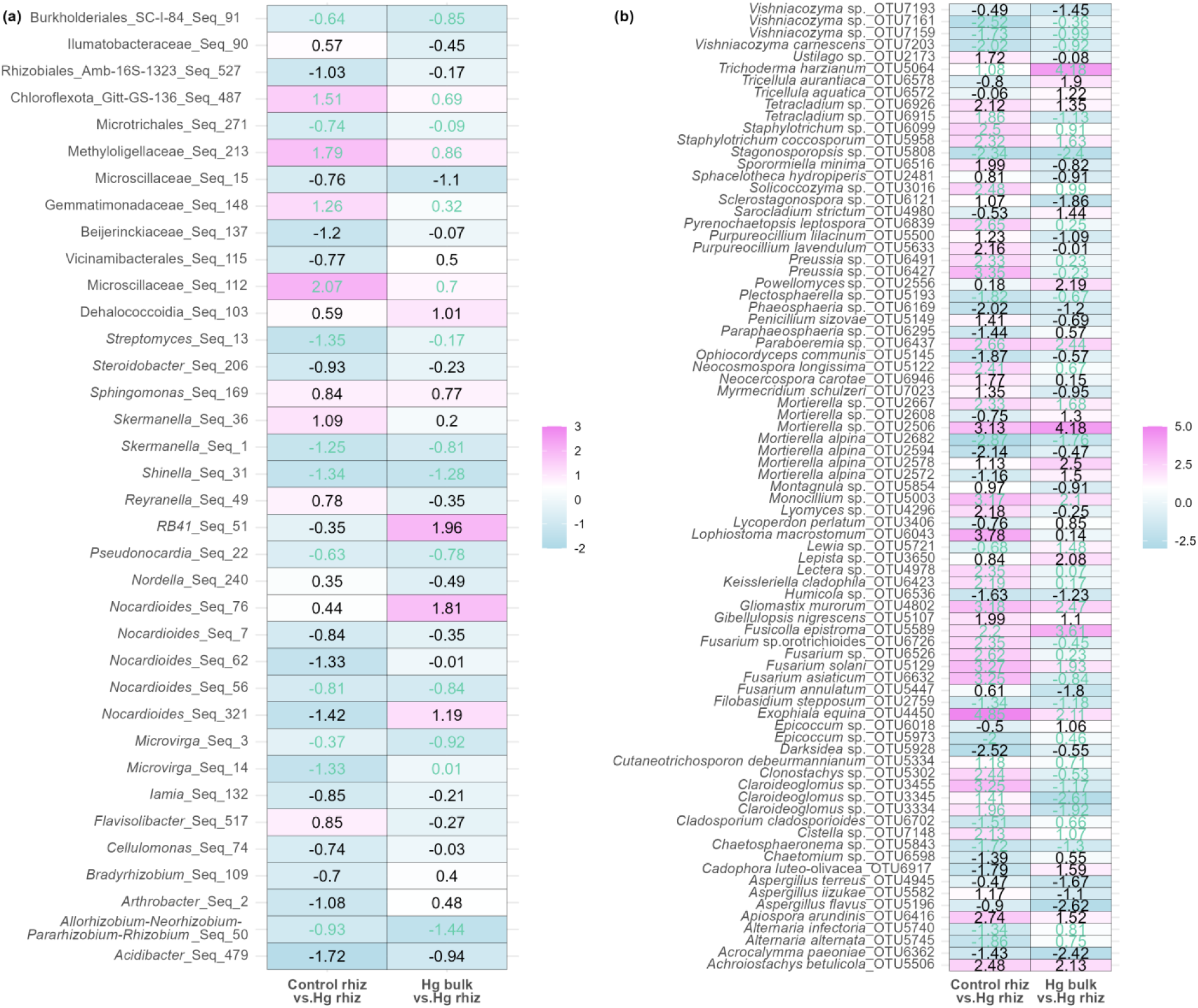
Heatmaps displaying differentially abundant genera/species in bacterial (a) and fungal (b) communities, comparing the Control rhizosphere (Control rhiz.) and Hg-contaminated bulk soil (Hg bulk) communities against the Hg-contaminated rhizosphere (Hg rhiz.). The ANCOM-BC2 global test was performed at the ASV (for bacteria) or OTU (for fungi) level. The log-fold change values for the globally significant taxa are shown inside each cell. Green entries denote taxa that have passed the sensitivity analysis for pseudo-count addition. The cells are color-coded to represent the direction and magnitude of log-fold changes (LFC): pink indicates an increase in abundance, while blue indicates a decrease for the microbial taxa in the compared group versus the Hg-contaminated rhizosphere. The Benjamini-Hochberg *p*-value correction method was applied to account for multiple testing. Unclassified ASVs were included in the analysis and figure with the closest identified higher taxonomic level.

Approximately 20% of the globally significant fungal taxa were differentially more abundant in Hg rhizosphere than Control rhizosphere and Hg bulk communities: *Vishniacozyma* sp. and *V. carnescens*, *Stagonosporopsis* sp., *Plectosphaerella* sp., *Ophiocordyceps communis*, *Mortierella alpina*, *Humicola* sp., *Filobasidium stepposum*, *Darksidea* sp., *Chaetosphaeronema* sp., *Aspergillus terreus*, *A*. *flavus*, *Acrocalymma paeoniae*.

Principal coordinates analyses were performed to determine patterns in the variation of the microbial communities structure. For the bacterial communities, 18.2% of the total variation in structure was explained by the principal axis of a constrained PCoA (CAP), which mainly separated the data points into clusters according to the sites of origin (Control vs. Hg-contaminated site) (Fig. 7a). This dissimilarity between sites was further affirmed with a significant (*p* < 0.001) PERMANOVA result (Table S6). The second constrained PCoA axis was responsible for explaining only 6.6% variation and was separating the data according to the Hg soil concentration. Moreover, the effect on bacterial community structure was also significant for soil Hg concentration across all sites (*p* < 0.01), sample type (bulk vs. rhizosphere soil) at the Hg site (*p* < 0.01), soil moisture (*p* < 0.001), and the presence/absence of the *merA* gene in the soil DNA (*p* < 0.001) (Table S6).

**Fig 7.**
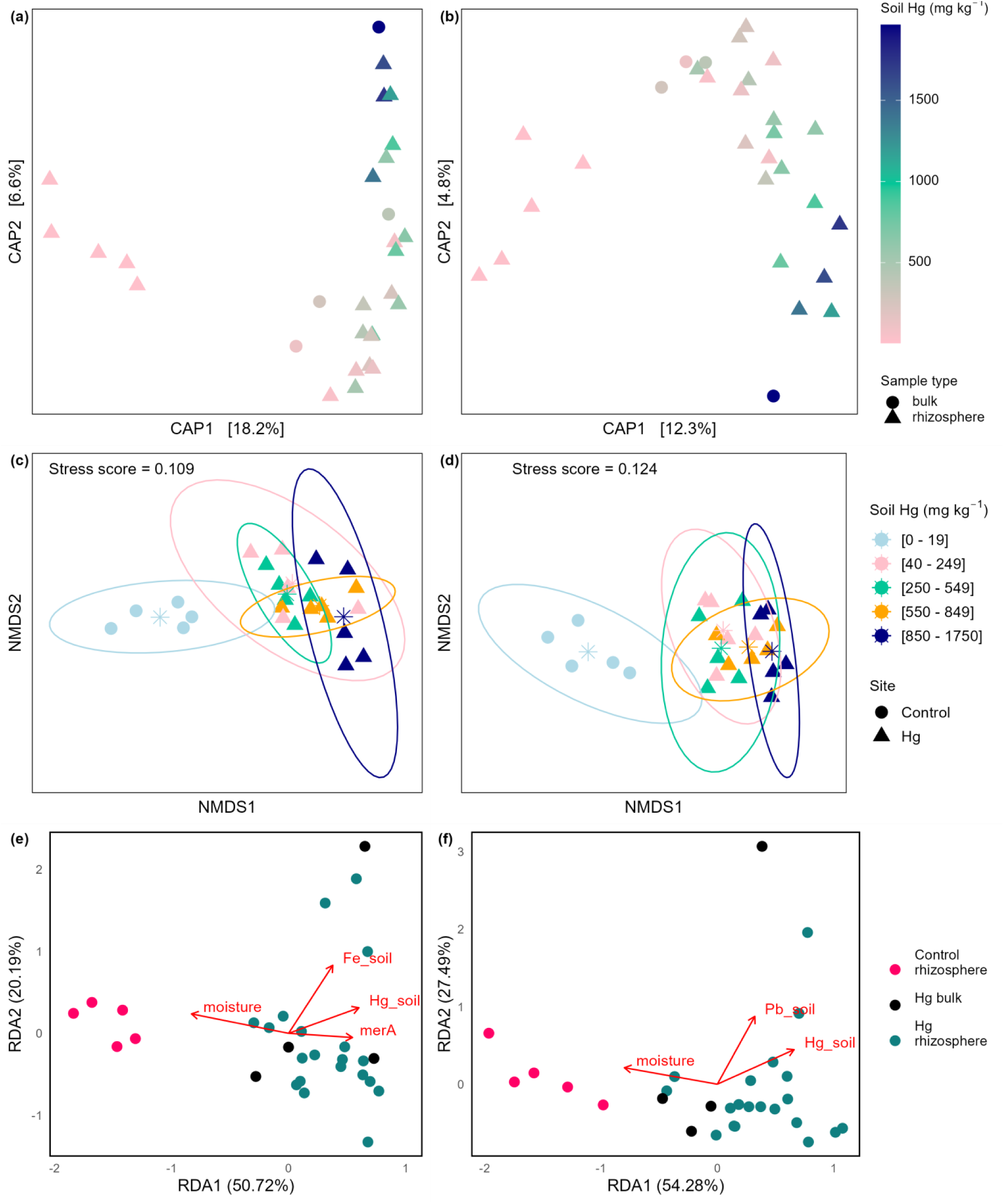
Principal coordinates analysis (PCoA) constrained by site and Hg soil concentration illustrating the relationships/ or differences between samples based on the Bray-Curtis dissimilarity matrices for the bacterial (a) and the fungal (b) communities in the bulk (full circles) and rhizosphere (full triangles) soil of *Lotus tenuis* at the Control and Hg-contaminated sites in Turda, Romania. Non-metric multidimensional scaling (NMDS) plots for the bacterial (c) and fungal (d) communities in the rhizosphere soil of *Lotus tenuis* at the Control and Hg-contaminated sites in Turda, Romania, depicting the dissimilarities in community composition according to Bray-Curtis distance matrices. The ellipses show the 95% confidence interval around the data points grouped according to the soil Hg concentrations. Distance-based Redundancy analysis (db-RDA) showing the total variation at the site explained by the environmental variable (red arrows) on the Bray-Curtis distance matrices of scores for the bacterial (e) and fungal (f) community structures at the Hg and Control sites.

The principal axis of the constrained PCoA on fungal communities explained 12.3% of the total variation and separated the communities into Control vs. Hg site although not as clearly as for the bacterial communities (Fig. 7b). The second PCoA axis separated the data according to the Hg soil gradient and explained 4.8% of variation. The PERMANOVA analysis revealed significant differences between communities’ structure based on site (*p* < 0.001) and soil Hg concentrations (*p* < 0.01) (Table S6).

Furthermore, for *L. tenuis* rhizosphere soil microbial communities, NMDS plots were generated after grouping the communities according to the level of Hg concentration in soil (Fig. 7c, d). Both bacterial and fungal communities clustered on the first axis according to the soil Hg level. However, significant differences in community structure were mainly found between each soil Hg concentration level at the contaminated site ([40-249 mg kg^-1^], [250-549mg kg^-1^], [550-849 mg kg^-1^], [850-1750 mg kg^-1^]) and the Control Hg level [0-19 mg kg^-1^], according to a pair-wise PERMANOVA (Table S7). The first two groups of soil Hg levels ([40-249 mg kg^-1^] and [250-549mg kg^-1^]) at the Hg-contaminated site were not clearly separated for neither bacterial nor fungal communities. There also seems to be less variation, less spread (lower confidence intervals) in the bacterial and especially fungal community structure of the Hg rhizosphere communities together than in the Control soil communities, based on the NMDS plots.

To determine the main environmental drivers of the community structure, redundancy analyses (RDA) were generated based on the Bray-Curtis distance matrices for bacterial and fungal community. The db-RDA models explained 31.5% and 20.9% of the total observed variation at the sites, for bacterial and fungal communities, respectively (ST8). The first axis of both db-RDA models explained around half of the constrained variation (Fig. 7e, f). For both models, the first axis was explained antagonistically by soil moisture and soil Hg concentration, and additionally for the bacterial communities the presence/ absence of the *merA* gene, which explained the highest proportion (ST8). Largely, Hg rhizosphere communities exhibited a similar trend for both bacteria and fungi, clustering on the first axis side driven by the *merA* presence and Hg soil concentration (bacteria) or Hg soil (fungi), whereas the Control communities are mainly driven by the moisture in soil.

### 3. Rhizosphere Microbial community and plant Hg-accumulation profile

The Hg-contaminated rhizosphere soil microbial communities were separated into groups based on the Hg plant shoot concentration, with low, medium, and high Hg shoot concentrations. Differentially more abundant ASVs/OTUs were revealed using the ANCOM-BC analysis. For bacterial communities no differentially more abundant species were detected in the rhizosphere of plants with high Hg shoot concentration compared to low and medium Hg concentrations (Fig. 8a). Fungal species more abundant in the rhizosphere soil of plants with high Hg shoot concentrations were *Septoglomus*, *Scytalidium*, *Plectosphaerella*, one OTU of *Mortierella alpina*, *Dioszegia*, *Articulospora*, *Alternaria*, and *Alfaria dandenongensis*.

**Fig. 8.**
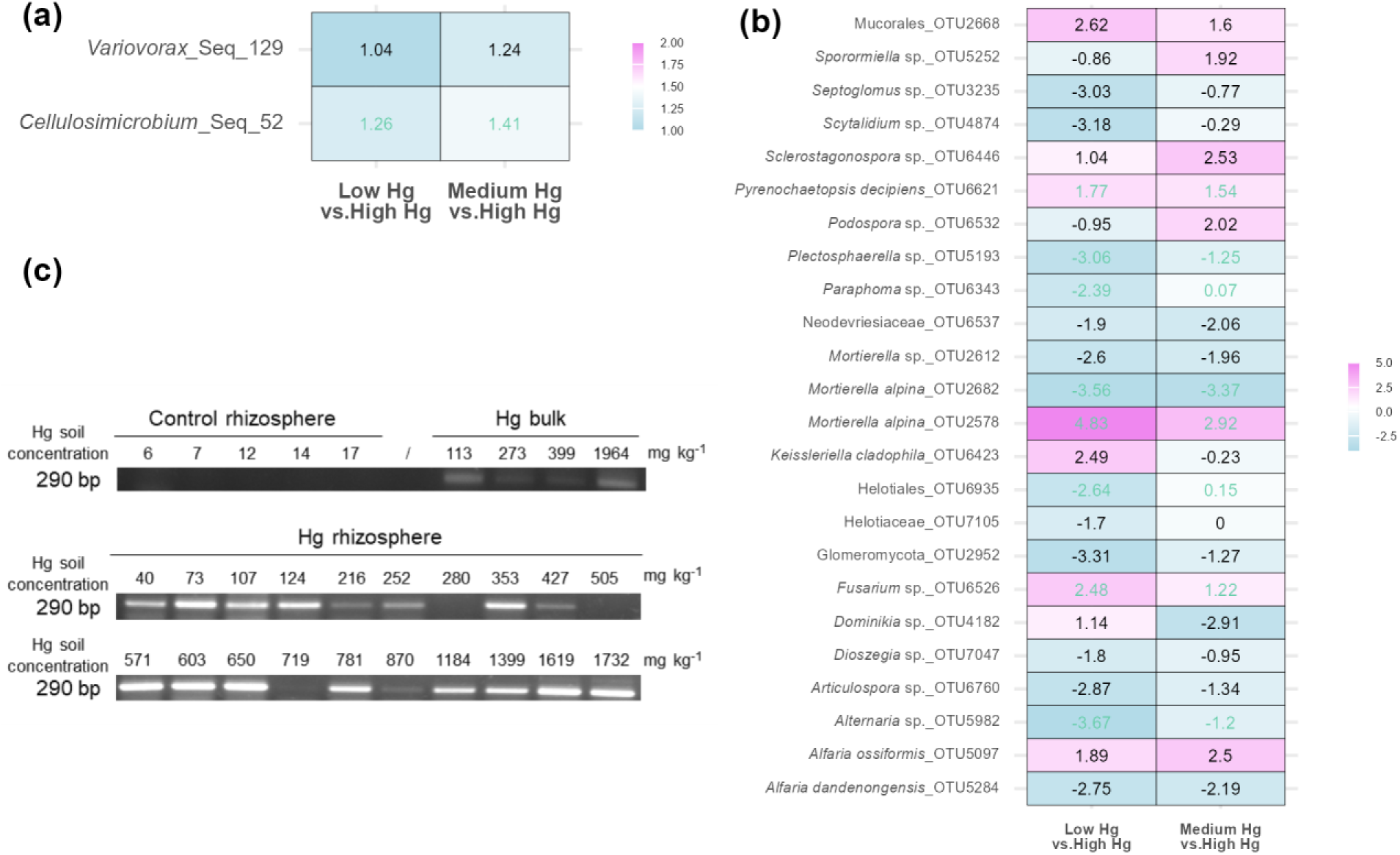
Heatmaps displaying differentially abundant genera/species in bacterial (a) and fungal (b) communities, comparing the rhizosphere soil communities of plants with low (Low Hg) and medium (Medium Hg) shoot Hg concentrations against communities from plant rhizospheres with high Hg concentrations in shoots (High Hg). The ANCOM-BC2 global test was performed at the ASV (for bacteria) or OTU (for fungi) level. The log-fold change values for the globally significant taxa are shown inside each cell. Green entries denote taxa that have passed the sensitivity analysis for pseudo-count addition. The cells are color-coded to represent the direction and magnitude of log-fold changes (LFC): pink indicates an increase in abundance, while blue indicates a decrease for the microbial taxa in the compared group versus the Hg-contaminated rhizosphere. The Benjamini-Hochberg *p*-value correction method was applied to account for multiple testing. *merA* gene amplicon presence (c) in total soil DNA samples from Control rhizosphere, Hg bulk, and Hg rhizosphere soil. Agarose gel electrophoresis, amplicon band at 290 bp length.

The potential mechanisms that would explain the community resilience under the Hg gradient was assessed by endpoint PCR, targeting the *merA* (mercuric reductase) gene, conducted on total DNA extracted from soil. 85% of Hg rhizosphere communities were positive for *merA*, as well as all Hg bulk communities (Fig.8c). As hypothesised, *merA* was not found in the Control rhizosphere soil communities.

### 4. Fungal classification into ecological guilds

A mere 15% of OTUs were matched to a representative species in the FungalTraits database and were assigned to an ecological guild. The plant pathogen guild was the most abundant per the three groups (Control rhizosphere, Hg rhizosphere, and Hg bulk) (Fig. S6). Ectomycorrhizal species were the second most abundant fungi, with the Hg rhizosphere communities having 4 times more than Control rhizosphere and 47 times more than Hg bulk. However, the relative abundance of the guilds varied greatly among samples; no trend could be observed as a response to the Hg gradient in the soil (ectomycorrhizal group - Pearson *r* = -0.06, *p* = 0.148).

## Discussion

Metal-contaminated sites pose long-term disruptions to ecological systems; however, they harbor uniquely adapted plant species and microorganisms. Assessing these environments can lead to the unexpected discovery of metal-tolerant or metal-avoidant plant species, as well as metal-resistant microorganisms, offering insights into adaptive mechanisms and potential applications in bioremediation (Reeves et al., 2018). Pioneer plant species developing advantageous genetic or epigenetic traits are valuable for plant-based remediation and management of contaminated soils (Krämer, 2010). Soil health depends on microbial processes (Schloter et al., 2018), with microbial community structure and function serving are indicators of soil type and quality (Nannipieri et al., 2017). The plant root rhizosphere, in particular, harbors a diverse and dynamic microbial niche driven by plant-secreted metabolites (Maurer et al., 2021). Thus, restoring metal-contaminated sites requires identifying the optimal holobiont—an integrated system of plants and their associated microbial communities.

The current study assessed the structure of the microbial communities inhabiting the rhizosphere of *L. tenuis* plants growing at an historically highly Hg-contaminated site. *Lotus tenuis* is a pioneer species, abundant at the investigated site, and also at sites contaminated with organic pollutants, such as hexachlorocyclohexane (HCH), highlighting its (epi)genetic potential of adaptability to abiotic stress (Balázs et al., 2018; Tiodar et al., 2024). To understand the impact of mercury contamination on plant rhizosphere microbial community dynamics, we compared Hg-tolerant rhizosphere soil microbial communities to those in non-contaminated rhizosphere soil and Hg-contaminated bulk soil. Significant shifts in the microbial community structure highlight the influence of multiple environmental and genetic factors. Key determinants of community dissimilarity included soil Hg concentration, the presence/absence of the plant, soil moisture, and presence/absence of the *merA* gene (for the bacterial communities), according to the significant PERMANOVA results.

Bacterial and fungal richness remained unaffected by the Hg soil gradient and the presence or absence of a host plant (Fig. 4). The resilience of bacterial community richness to Hg contamination was likely due to the widespread presence and potential activity of Hg detoxification systems, as indicated by the high presence of the *merA* gene in all rhizosphere soil samples (Fig. 8c). This contrasts with the response of rhizosphere communities of the same plant species under HCH stress, where microbial richness declined with increasing soil HCH concentration, highlighting distinct adaptive capacities to different contaminants (Balázs et al., 2021).

The resilience of bulk soil microbial richness under Hg stress has been previously observed, with bacterial and fungal communities maintaining diversity even in soils contaminated with 155, 840, and 1,130 mg kg⁻¹ Hg and high arsenic concentrations (Prosenkov et al., 2023). However, at an Hg-contaminated site undergoing poplar-assisted phytomanagement, fungal richness significantly declined along a gradient of 3–9 mg Hg kg⁻¹ soil (Durand et al., 2017). These contrasting findings may stem from differences in the nature of Hg stress: phytomanagement strategies could create fluctuations in soil Hg availability, whereas chronic Hg contamination, as in the present study, establishes a more stable gradient. Alternatively, differences in Hg concentration levels could explain the discrepancy. If so, fungal species richness may stabilize under prolonged exposure to high Hg concentrations (40–1,964 mg kg⁻¹) or at the levels tested by Prosenkov et al. (2023). In this scenario, only highly adapted Hg-resistant fungal species persist, developing facilitative interactions that mitigate Hg toxicity and enable sensitive species to regain abundance. Conversely, under lower Hg stress gradients, competition from more tolerant strains may drive sensitive taxa to decline as Hg concentration increases.

Similar to richness, bacterial alpha diversity in Hg-contaminated rhizosphere communities remained largely unchanged compared to the Control site. However, the relationship between bacterial alpha diversity and soil Hg concentration has yielded contrasting findings across studies. Frossard et al. (2018) reported an increase in community richness and diversity in Hg-tolerant bulk soil communities exposed to higher Hg concentrations compared to those from lower-Hg areas. In contrast, Liu et al. (2014) observed that alpha diversity in paddy bulk soil communities plateaued at 5 mg Hg kg⁻¹ within an Hg gradient of 1–29 mg kg⁻¹. Similarly, Prosenkov et al. (2023) found no significant differences in bacterial and fungal community diversity across three high-Hg soil concentrations. These findings suggest that in the present study, rhizosphere bacterial communities might have reached an alpha diversity plateau within the Hg stress gradient of 40–1,732 mg kg⁻¹, likely reflecting a threshold beyond which diversity remains stable despite increasing contamination.

However, in this study, the fungal communities in the Hg rhizosphere were statistically the least diverse compared to the Control or Hg bulk soil, suggesting that plants select for specific taxa in the Hg-contaminated environment. The dominance of Ascomycota was confirmed for the Hg rhizosphere communities, as it is for most soil habitats (Egidi et al., 2019). Similarly, Ascomycota and Basidiomycota dominance was observed by Frossard et al., 2018, in a bulk soil chronically contaminated with up to 36 mg Hg kg^-1^.

At the bacterial phylum level, the relative abundance of Gemmatimonadota was of interest, as it presented a linear, albeit weak relationship (R^2^=0.26, *p*<0.01, Supplementary Fig. 6) with Hg soil concentration. This trend was previously confirmed with a significant and strong linear dependency of Gemmatimonadota to the chronically Hg concentrations in bulk soil following a gradient of 1-29 mg kg^-1^ in the study of Liu et al., 2014. The decrease in the strength of the relationship could be explained by the increased soil Hg concentrations of the gradient in our study, signifying that Gemmatimonadota might be an indicator of Hg-contamination in the lower to moderate concentration ranges. Moreover, the difference between our study and Liu et al could also be influenced by the soil pH, as for our study the pH measured in CaCl_2_ could indicate a more alkaline pH compared to the soil pH in the study by Liu et al.

In this study, Bacillota exhibited a significant negative linear relationship with soil Hg concentration (R² = 0.39, *p* < 0.001, Fig. S7), with a sharper decline from the control communities toward the Hg-contaminated soil. In contrast, Liu et al. (2018) reported a weak but significant positive correlation between Bacillota abundance and total Hg concentration in bulk soil bacterial communities from chronically Hg-contaminated upland farmland adjacent to paddy soils, across an Hg gradient of 0.3–52 mg kg⁻¹. Furthermore, in this study, the genus *Clostridium* sensu stricto 1 within Bacillota showed no clear relationship with soil Hg concentration based on relative abundance (R² = 0, *p* = 0.864, Fig. S8), despite being three times more abundant at the Hg-contaminated site than at the Control site. In contrast, Liu et al. (2018) observed a significant positive correlation between *Clostridium* and Hg concentration. These conflicting patterns may be attributed to differences in Hg gradients between studies, as well as differences in sampling environments—Liu et al. analyzed bulk soil, whereas the present study focused on rhizosphere soil from a pioneer plant, where plant-microbe interactions and root exudates may have influenced microbial community responses to Hg contamination.

A few bacterial orders were highly abundant across the 16S dataset, and had similarly high relative abundance in all communities, Hg rhizosphere, Hg bulk, and Control samples. Solirubrobacteriales, Propionibacteriales, and Micrococcales, all from the Actinomycetota phylum were the common soil microorganisms adapted to the climatic conditions of the two sites exhibiting also resilience to edaphic changes. The two sites are in close proximity and share the same temperate continental climate. However, the Hg site is an industrially used brownfield with Orthent soil dominated by sandy texture and low in organic material, therefore with low water retention properties, prone to erosion with a semi-arid degraded landscape, especially during the summer months (and at the time of sampling). Despite these environmental differences at the two sites, the pH was neutral to slightly alkaline for both, explaining the resilience of Actinomycetota members as they grow optimally in this pH range (Lauber et al., 2009).

The orders Sphingomonadales (Alphaproteobacteria) and Burkholderiales (Betaproteobacteria) were highly abundant in Hg-contaminated rhizosphere and bulk soil, but not in Control soil, with Sphingomonadales exhibiting the highest abundance in bulk soil. This trend was maintained for Sphingomonadaceae, the fourth most abundant bacterial family across the dataset, up to its representative *Sphingomonas* member, that was most abundant in Hg bulk soil and decreased 2-fold in Hg rhizosphere and 4-fold in Control rhizosphere soil (Supplementary Table 8). However, Burkholderiaceae was not among the most abundant families, and was solely represented by *Lautropia* and *Limnobacter*, both more abundant at the Hg site than at the Control (Supplementary Table 8). These results suggest that both families are not part of the core *L. tenuis* rhizomicrobiota, but the plant root acquisition of these taxa might have occurred later at the Hg-contaminated soil. These results are in contrast to the study by Luo et al., 2022, which found that the core rhizosphere of 5 Cd-(hyper)accumulator plants were enriched in ASVs of Burkholderiaceae and *Sphingomonas* compared to the bulk soil of the rhizobox where the plants were cultivated under high and low Cd-contaminated and non-contaminated soil conditions. In our study, *Burkholderia* was not present among the identified ASVs. *Lotus tenuis* is a Hg-indicator plant species (Tiodar et al., 2024) that has exhibited tolerance to HCH as well (Balázs et al., 2018) and might thus employ distinct strategies of acquiring beneficial/ core rhizo-microbiota compared to metal accumulator species that consistently exhibit preference to one or two metal species. Under multiple stress factors, such as high Hg and aridity, *L. tenuis* might favor a more diverse rhizo-microbiota rather than a sharp one, enabling it to cope with the complexity of stressors.

To identify potential microbial partners that mitigate the stress of Hg in soil, the abundance of bacterial and fungal taxa was compared (ANCOM-BC) between sites. A few ASV-level taxa were differentially more abundant in the rhizosphere at the Hg-contaminated site than at the Control site, or in the bulk soil. Bacterial taxa found in the rhizosphere soil of the Hg tolerant ecotype were *Streptomyces*, *Nocardioides, Skermanella*, *Steroidobacter*, *Shinella*, *and Allorhizobium-Neorhizobium-Pararhizobium-Rhizobium*.

Members of the *Streptomyces* genera (Actinomycetota) are generally plant beneficial symbionts that can potentially promote plant growth (Olanrewaju & Babalola, 2019) by solubilizing the pool of soil phosphorus into polyphosphate, the form available for plant uptake (Alori et al., 2017). Moreover, *Streptomyces* sp. can alleviate biotic and abiotic stress, in the rhizosphere functioning as a biocontrol agent promoting plant health through the extensive production of bioactive secondary metabolites, not only antibiotics, but also pesticides and herbicides (Ankati et al., 2021; Pang et al., 2022). A few metal-resistant strains of *Streptomyces* have also been isolated (Al-Huqail & El-Bondkly, 2022), and *S. pactum* has been used for improving the soil metal (Cd, Cu, Pb, Zn) remediation traits of *Triticum aestivum* in a pot experiment with promising benefits such as 32% shoot biomass increase and 2-fold increased Cd shoot accumulation in inoculated wheat plants compared to uninoculated plants under a Cd stress of 118 mg kg^-1^ soil (Ali et al., 2021). In the current study, a reduction of pathogenic fungal species in the rhizosphere compared to bulk soil was noticed for species such as *Fusarium solani, Alternaria infectoria* and *A. alternata* at the Hg site (Fig. 6b.) suggesting an inter-kingdom community shift towards plant health promoting microbiota in the rhizosphere. This dynamic trend of restructuring the community towards plant positive microbiota, was also observed in the study of Feng et al., 2024, where *Streptomyces* together with *Sphingomonas* were enriched concurrently with *Fusarium* in the root endophyte microbiota of maize plants exposed to 20 mg Hg kg^-1^ soil. It was thus hypothesized that the increase in relative abundance of *Streptomyces* and *Sphingomonas* might have had a *Fusarium* bio-stress alleviating effect on maize plants. Interestingly, in this study the mean relative abundance of *Fusarium* species was maintained at the same threshold in Hg rhizosphere soil as in bulk soil, and twice lower than in Control rhizosphere soil. These results suggest that *L. tenuis* has adapted to chronic Hg stress by restructuring the rhizosphere microbiota to maintain the abundance of *Fusarium* species under a disease-inducing level.

*Nocardioides*, another Actinomycetota genus abundant in the Hg-contaminated rizosphere has a wide distribution due to its adaptation to low-nutrient conditions and pollutant degradation potential, especially for organic classes (Y. Ma et al., 2023). It also has the potential to acquire metal-resistance properties, as was seen for Cu (Xavier et al., 2019). Moreover, this is not the first account of *Nocardioides* being abundant in a Hg-contaminated soil (D. Li et al., 2022).

*Skermanella* of the Azospirillaceae family is a genus typically found in soil in a variety of habitats from meadow soils (X. Guo et al., 2020) to desert sands (Subhash & Lee, 2016b). Members of *Skermanella* are unable to fix C or N (Zhu et al., 2014). Moreover, a strain of *S. stibiiresistens* sp. nov. was isolated from contaminated coal mine soil and proved to be highly resistant to As (Luo et al., 2012) suggesting potential in acquiring resistance against toxic metals. The role of the Gammaproteobacteria of the *Steroidobacter* genera is currently undetermined. However, it seems to be restricted to the rhizosphere area as some strains (S. *agaridevorans* sp. nov.) require metabolites for growth that are regularly secreted by members of the Rhizobiales order (Ikenaga et al., 2021). In this study, two genera of the Rhizobiaceae family, *Shinella* and *Allorhizobium-Neorhizobium-Pararhizobium-Rhizobium* were differentially more abundant in the Hg-contaminated rhizosphere than in Control rhizosphere or Hg-contaminated bulk soil. Although *Shinella* species usually test negative for nitrogen fixation (An et al., 2006), there has been an instance of *S. kummerowiae* sp. nov. being positive for the presence of the *nifH* gene when isolated from the root nodules of a legume plant (*Kummerowia stipulacea*; Lin et al., 2008). Similarly to *Skermanella*, *Shinella* strains (*S. curvata* sp. nov.) have been isolated from hydrocarbon-contaminated desert sand (Subhash & Lee, 2016a) suggesting an adaptation potential to water depleted and contaminated soils. Although *Allorhizobium, Neorhizobium, Pararhizobium*, and *Rhizobium* can be distinguished as individual genera, the SILVA taxonomy classifies them as one genus. As such*, Allorhizobium-Neorhizobium-Pararhizobium-Rhizobium* bacteria are usually found in root nodules and can convert atmospheric nitrogen into ammonium nitrogen, and thereby increase the pool of plant-available N in the rhizosphere of the Hg-contaminated soil enhancing the plant’s nutritional status (Velázquez et al., 2017).

The rhizosphere microbiota of tolerant plants growing in contaminated areas is also important due to the organic decomposers (*Streptomyces*, *Pseudonocardia*, *Cellulomonas* etc.) that fertilize the soil and prepare the advance of less tolerant plant species that require higher nutrient content. Albeit not differentially more abundant in the Hg-contaminated rhizosphere compared to Control and bulk soil, it is worth mentioning that *Mesorhizobium* and *Bradyrhizobium* had high relative abundance at both sites, both genera containing nitrogen-fixing and nodule-forming members (Supplementary Table 8). *Mesorhizobium* is a recognized nodule symbiont of *Lotus* species (Crosbie et al., 2022) and herein has a high relative abundance in the rhizosphere of both Hg-contaminated and Control soils, being 4 times less abundant in the Hg-contaminated bulk soil compared to rhizosphere soils. The specific symbiont of *Lotus tenuis* was previously identified as *Mesorhizobium* and it is considered that *M. loti* is the rhizobia symbiont inducing nodule formation in *Lotus* species from the *L. corniculatus* clade (Lorite et al., 2018; Saeki & Kouchi, 2000). However, *Bradyrhizobium* had similar mean relative abundance in all three soil categories, while being 2-fold less abundant than *Mesorhizobium* in rhizosphere samples reinforcing the knowledge that *L. tenuis* is not its preferred host (Jarvis et al., 1982). Nonetheless, the presence of *Bradyrhizobium* species at Hg-contaminated sites could be a favorable setting for future crop plants to establish at the site. For example, soybean is known to form symbiosis with particular species of *Bradyrhizobium* which proliferate in nodules even when their abundance in bulk or rhizosphere soil is limited (Sarao et al., 2024).

Furthermore, *Pseudomonas* was prevalent in the soil at the Hg site. In this study, the mean relative abundance of *Pseudomonas* was 4 times higher at the Hg site compared to the Control site. This is not surprising as *Pseudomonas* is a known soil bacterium with Hg-resistance traits, as assessed by numerous studies (Bourdineaud et al., 2020; Ginting et al., 2021; Holtze et al., 2006; Ji et al., 2018; Lo et al., 2022; Oliveira et al., 2010; Uqab et al., 2024). For example, Zappelini et al., 2015 discovered a 19-fold increase in the total number of *Pseudomonas* in soil disturbed by chlor-alkali tailings dump enriched in Hg and As, compared to undisturbed soil. However, *Pseudomonas* was also 2 times more abundant in rhizosphere than in bulk soil. For the nodule microbiota of *Lotus* species, *Pseudomonas* is an important non-rhizobia symbiont, that can establish positive interactions with effective nodulating rhizobia species (e.g. *Mesorhizobium*), while through negative microbe-microbe interactions diminishes the risk of ineffective nodules formed by other rhizobia species (e.g. *Allorhizobium-Neorhizobium-Pararhizobium-Rhizobium*) (Crosbie et al., 2022). This is especially important, as herein both *Mesorhizobium* and *Allorhizobium-Neorhizobium-Pararhizobium-Rhizobium* are in high abundance in the Hg-contaminated rhizosphere soil and thus the risk for the plant to form costly ineffective nodules is high.

Our study revealed notable differences in the bacterial community composition at the genus level in Hg-contaminated soil from previous works such as by Ma et al., 2019, which examined rice rhizosphere and bulk soil communities under a soil Hg gradient. The 10 most abundant genera reported in Ma et al. are distinct from the most abundant ones found in this study at the Hg site based on mean relative abundance. Out of the three genera that were enriched in the high Hg sites in their study, only *Xanthomonas* overlapped with our dataset, and herein it was completely restricted to 35% of the Hg rhizosphere communities. However, different taxonomic profiles between the two studies are not surprising, since the two soil Hg gradients are not coinciding ([0-5] (Ma et al., 2018) vs. [30-2000] mg Hg kg^-1^ soil).

Noteworthy is also the presence of *Microbacterium* sp. (Actinomycetota) restricted to the Hg-contaminated site. Mariano et al., 2020 identified *M. trichothecenolyticum* as the most abundant Hg indicator species among the cultivable rhizosphere bacteria of two plant species (*Aeschynomene fluminensis* and *Polygonum acuminatum*) colonizing a site contaminated with 3 mg Hg kg^-1^ soil. In our study, soil Hg concentration was 10 to 667 times higher, and *Microbacterium* had a mean relative abundance in rhizosphere twice higher than in bulk soil. The *M. trichothecenolyticum* strain isolated by Mariano et al. was positive for *merA* presence and for a few plant growth-promoting properties (secretion of siderophores, proteases, and indoleacetic acid synthesis), which would explain its presence in Hg-contaminated substrates (*merA*) and increased abundance in rhizosphere soil.

It appears that plant roots at the Hg-contaminated site influence the rhizosphere bacterial community, selecting microbes that provide a combination of plant growth promoting traits, nutrient acquisition, pathogen suppression, aridity (a co-stress factor at the site), and Hg tolerance. Hg-tolerant bacteria likely play a critical role in supporting the microbial community by reducing Hg toxicity through mechanisms such as the mercuric reductase system, which alleviates stress for neighboring microbes. Thus, the results of this study suggest that plant-mediated selection in the rhizosphere prioritizes bacterial traits beneficial to both plant health and microbial community resilience, particularly from the pool of *merA*-harboring bacteria. This is consistent with a recent metagenomic study from a Hg-contaminated site in Almadén, Spain (González et al., 2022), which identified *merA* resistance mechanisms in both bulk and rhizosphere soils but reported a higher abundance of plant-beneficial traits and stress-response genes, such as those related to salinity and desiccation, in the rhizosphere. Therefore, the results of this study indicate that the stability of the bacterial rhizosphere community under the Hg stress gradient is likely driven by cooperative and complementary interactions among diverse microbes with multifunctional roles.

Fungal communities were less diverse in the Hg-contaminated *L. tenuis* rhizosphere soil than in bulk or Control rhizosphere soil, in terms of low alpha diversity. This community trend was confirmed at the order level, with Pleosporales (Ascomycota) being clearly dominant at the Hg site, especially in the rhizosphere communities, with a mean relative abundance of 25%, 3 times higher than in the Control rhizosphere. This is not surprising, as previously the ecologically diverse Pleosporales was found dominant in semiarid landscapes, aridity being an additional detrimental condition at the Hg site (Knapp et al., 2015; Ndinga-Muniania et al., 2021; Porras-Alfaro et al., 2011).

Interestingly, the Agaricales (Basidiomycota) order was abundant in rhizosphere soil at both the Hg-contaminated and Control sites, in almost equal proportions, but 4 and 3 times respectively less abundant in the Hg bulk soil, suggesting a shared presence in the core rhizofungal communities of *L. tenuis* and resilience to Hg stress. In contrast, Glomerales (Mucoromycota), Pezizales (Ascomycota), and Filobasidiales (Bazidiomycota) exhibited preference for the rhizosphere of the Hg-contaminated soil, as for each the relative abundance was at least a 2- to 3-fold increased therein compared to both the Control rhizosphere and bulk soil. This trend suggests these fungal orders may have members exhibiting ecological adaptations, such as Hg tolerance that allow them to thrive in the contaminated soil and plant beneficial traits that favor their proliferation in the rhizosphere niche. Arbuscular mycorrhizal (AM) or ectomycorrhizal symbionts of Glomerales and Pezizales respectively could establish mutually beneficial relationships with plants in exchange for plant-secreted nutrients (Bago et al., 2000; Tedersoo et al., 2006). Glomerales is an important order of the Glomeromycotina subphylum known for containing almost exclusively fungi that can establish arbuscular mycorrhizal symbiosis with many land plants, including legumes (Parniske, 2008). The multi-partite legume symbiosis model validated also for *L. japonicus*, a close relative of *L. tenuis*, is the basis of the pioneer functionality of legumes in nutrient-depleted and contaminated soils (Handa et al., 2015; Tsiknia et al., 2021). By initiating an additional symbiosis with fungi, legumes extend their nutrient-acquiring networks to include fungi as well as bacteria, and their microbe-acquired nutrients repertoire to include phosphate (Harrison et al., 2002). It is likely that AM fungi are of critical importance for the *L. tenuis* at the Turda site not only for acquiring minerals but especially water (Kakouridis et al., 2022). Kakouridis et al. previously demonstrated water transfer to host plants using *Rhizophagus intraradices* (Glomerales) as a model. Indeed, herein, *Rhizophagus* sp. was three times more abundant in the rhizosphere of the Hg-contaminated soil than in the Control soil (Supplementary Table 8). This aligns with the observation that soil moisture content in the Hg-contaminated rhizosphere was three times lower than in the Control site, yet the shoot moisture content was statistically similar for plants at both sites (Fig. S1e).

Pezizales was thought to consist mainly of saprotrophic species, but eventually it was shown to include a large representation of ectomycorrhizal fungal lineages (Tedersoo et al., 2009). Interestingly, in this study, an abundant pezizalean genus was *Peziza*, which has a few ectomycorrizal members (Tedersoo et al., 2006). Moreover, *Peziza* sp. was highly abundant at the Hg rhizosphere soil, while virtually missing in bulk soil communities and in the Control rhizosphere soil (Supplementary Table 8). In the study by Durand et al., 2017, *Peziza* sp. was the keystone species in the fungal network of Hg-contaminated poplar phytomanaged field for the soil, root, stem, and leaves fungal communities. These findings underscore an adaptive potential to Hg-induced stress and a preference for the rhizosphere habitat driven by the ectomycorrhizal-plant host trade-off mechanism.

Furthermore, the abundance of symbiotic fungal associations was also suggested by the outcome of the FungalTraits analysis. Albeit the fungal guild classification was limited to only 15% of total fungal OTUs, the Hg-contaminated rhizosphere appeared more abundant in beneficial interactions between plant roots and fungi, as more ectomycorrhizal species were found in Hg-contaminated rhizosphere compared to Control rhizosphere or bulk soil communities.

Among the differentially more abundant fungal taxa in the Hg-contaminated rizosphere soil were members of Pleosporales (*Stagonosporopsis* sp., *Darksidea* sp.*, Chaetosphaeronema* sp., *Acrocalymma paeoniae*), Glomerellales (*Plectosphaerella* sp.), Mortierellales (*Mortierella alpina*), Filobasidiales (*Filobasidium stepposum*), Sordariales (*Humicola* sp.), Tremellales (*Vishniacozyma* sp. and *V. carnescens*), and Eurotiales (*Aspergillus terreus*, *A*. *flavus*).

*Stagonosporopsis* species are generally identified as plant pathogens and saprotrophs, although recently new species were isolated from garden soil (Hou et al., 2020) and rhizosphere communities of certain plants, such as poplar (Wei et al., 2021). Previously, a *Stagonosporopsis* sp. P2.12 isolate, from the investigated site, displayed in culture very high and moderate resistance to 518 mg kg^-1^ Pb and 10 mg kg^-1^ Hg, respectively (Văcar et al., 2021).

*Darksidea* species are part of the dark septate root endophytes which is a group of plant beneficial fungi that promote plant growth and tolerance to abiotic stresses, including metal stress (Zhang et al., 2008) and aridity (Knapp et al., 2012, 2015; Li et al., 2018). *Acrocalymma paeoniae* is another dark septate fungus. Although studies on *A. paeoniae* are lacking, there are indications that *A. vagum* assists the growth of tobacco plants when cultivated under 5 Cd mg kg^-1^ soil pots, by promoting the development of a higher number of leaves and wider leaf area than uninoculated plants, and by reducing the Cd content in leaves up to 19-26% (Jin et al., 2018).

Another abundant pleosporalean genus in the Hg tolerant communities was *Chaetosphaeronema*. Members of this genus could have plant growth-promoting properties, such as *Chaetosphaeronema achilleae*. In a pot experiment, wheat plants inoculated with *C. achilleae* isolated from the rhizosphere of a plant native to arid soils (*Peganum harmala*) had significantly higher shoot dry biomass than uninoculated wheat plants (Mohamed et al., 2022). Moreover, extracts of plant associated *C. achilleae* exhibited antimicrobial properties against a few human pathogens (Narmani et al., 2019).

*Mortierella alpina* is an oleaginous endophyte known for its ability to enhance plant growth. In a study by Wani et al., 2017, inoculation with *M. alpina* significantly promoted the growth of *Crocus sativus*, leading to increased total biomass, larger corm size, and a greater number of adventitious roots compared to uninoculated controls. Moreover, *M. alpina* exhibits biocontrol properties against soil-borne *Fusarium oxysporum*, as demonstrated in a study on ginseng cultivated in soil pot experiments (Y. Wang et al., 2022). Not only inoculation with *M. alpina* inhibited the symptoms of ginseng root rot and decreased the relative abundance of *F. oxysporum* compared to non-inoculated plants, in the study by Wang et al., 2022, but plant growth was also promoted potentially through its indoleacetic acid production properties and enhanced availability of N and P. Furthermore, in the same study, *M. alpina* altered the architecture of rhizosphere soil communities by stimulating the ginseng recruitment of plant-beneficial microbes (*Pseudomonas*, *Sphingomonas*, *Rhizobium*). However, the potential beneficial effects of *M. alpina* might be species-specific, and therefore *in vitro* studies are necessary to elucidate the relationship with different host plants, such as *L. tenuis*. Moreover, this fungus was frequently isolated from the investigated site, displaying moderate to high resistance to 490 mg kg^-1^ Zn in culture (Văcar et al., 2021).

The *Vishniacozyma* genus identified as differentially more abundant in the Hg rhizosphere soil includes plant endophytes that have been previously investigated for their biocontrol agent properties against phytopathogenic fungi (e.g. *V. carnescens* (syn. *Cryptococcus carnescens*) against Dutch elm disease pathogen, *V. victoriae* (syn. *Cryptococcus victoriae*) against *Botrytis cinerea, Penicillium expansum*, *Cladosporium* sp., *Phlyctema vagabunda;* (Gorordo et al., 2022; Gramisci et al., 2018; Marčiulynas et al., 2022; Nian et al., 2023; Sepúlveda et al., 2022).

The *Humicola* genus holds some potential as a beneficial plant rhizosphere fungus. Strains of *Humicola* sp. and *Humicola phialophoroides* were previously identified in soils with plant pathogen suppressing properties (Ko et al., 2011; Lang et al., 2012).

The filamentous fungus, *Aspergillus terreus* is a functionally diverse organism that can be both a pathogen to plants and humans, a source of secondary metabolites with pharmaceutical properties, and a biocontrol agent against certain plant pathogens (Abdelaziz et al., 2023; Halo et al., 2018; X. Huang et al., 2021; Louis et al., 2014). *Aspergillus flavus*, a saprophytic fungus, on the other hand, has potent pathogenic properties to both plants and humans, and no observable positive effects on plant growth. It infects seed crops with the aflatoxin metabolite which causes the aspergillosis disease in humans (Amaike & Keller, 2011; Z. Huang et al., 2023; Klich, 2007). Nevertheless, *Aspergillus* spp. have been proposed and studied for their heavy metal remediation potential (Oladipo et al., 2016; L. Wang et al., 2024), while two *Aspergillus* sp. isolates from the investigated site exhibited high resistance in culture at 10 mg kg^-1^ Hg (Văcar et al., 2021). Thus, it is likely that *Aspergillus* spp. occupy an engineering niche within *L. tenuis* rhizosphere, achieving metals immobilization.

Species of *Plectosphaerella,* such as *P. citrulli*, *P. pauciseptata*, and *P. ramiseptata,* are plant pathogens (Raimondo & Carlucci, 2018). In this study, *Plectosphaerella* was more abundant in the rhizosphere of plants with high Hg concentrations in shoots. However, neither high Hg concentration in shoots nor the abundance of *Plectosphaerella* decreased the plant shoot DW at the Hg-contaminated site (Fig. S9).

In summary, the soil fungal network of the Hg-contaminated *L. tenuis* rhizosphere traded diversity to some extent in favor of highly selected fungal partners, with a preference for arbuscular/ectomycorrhizal symbionts. This enrichment in nutrient-cycling, mutually beneficial fungi and bacteria is a key sustainable advantage of colonizing contaminated sites with pioneer legume species. The abundant fungal genera were selected for their adaptability to abiotic stress, especially aridity and metal contamination. However, due to the extended presence of bacterial *merA* genes in the Hg-contaminated rhizosphere soil, it is herein hypothesized that bacteria are the main Hg-resistance line, with secondary support from the fungal side, especially the phytopathogenic species present in abundance in rhizosphere soil, whereas most fungal members better assist in supplying plant roots with their water and macronutrient requirements. Nonetheless, this fungal network also exhibits pathogen suppressing characteristics which could be useful in counteracting the invasion of opportunistic potentially Hg-resistant pathogens (e.g. *Aspergillus flavus, Plectosphaerella* sp., *Fusarium* sp.) that also inhabit the *L. tenuis* rhizosphere. Nonetheless, the fungal species more abundant in the rhizosphere soil of Hg-tolerant plants that accumulated the highest Hg concentration in shoots are of interest for the selection of future phytoremediation efficient microbial consortia. Noteworthy are the *Septoglomus*, with arbuscular mycorrhizal members (Błaszkowski et al., 2014; Guillén et al., 2020), *M. alpina*, which was also differentially more abundant in the rhizosphere at the Hg site than at the Control site or in the bulk soil, *Dioszegia*, which might play a crucial role in the diversity and stability of the plant phyllosphere microbiota (Agler et al., 2016), and *Articulospora*.

## Conclusions

The rhizosphere microbial communities of *Lotus tenuis* in Hg-contaminated soils exhibit remarkable functional resilience, maintaining stable bacterial and fungal diversity across an extensive Hg gradient (40–1964 mg Hg kg⁻¹ soil). Despite significant shifts in community structure between Hg-contaminated and control rhizosphere soils, alpha diversity remained largely unchanged, likely due to the widespread presence of the *merA* gene (85% of Hg-contaminated rhizospheres), highlighting bacterial detoxification as a key adaptive strategy. These findings support the stress gradient hypothesis, suggesting that facilitative interactions drive microbial adaptation under Hg stress rather than competition-driven diversity loss. At the Hg-contaminated site, the rhizosphere was enriched with *Mesorhizobium* sp., a core N-fixing symbiont, and *Pseudomonas* sp., a promising Hg-resistant bacterium with potential rhizosphere interactions optimizing nodulation. Low soil moisture likely drove plant selection of arbuscular mycorrhizal fungi, particularly *Rhizophagus*, to enhance water uptake. Several bacterial taxa, including *Streptomyces* sp. (a biocontrol agent), *Allorhizobium-Neorhizobium-Pararhizobium-Rhizobium* sp., and *Shinella* sp. (plant-growth-promoting rhizobia), as well as the less-studied *Nocardioides* and *Skermanella* (potentially metal- and aridity-resistant), were significantly enriched in Hg-contaminated rhizospheres. Key fungal partners, such as *Darksidea* sp. and *Acrocalymma paeoniae* (dark septate endophytes), *Mortierella alpina* and *Chaetosphaeronema* (potential plant-growth promoters), and *Humicola* sp. and *Vishniacozyma* sp. (pathogen suppressors), contribute to the functional resilience of the microbial network. This study provides a unique perspective by analyzing rhizosphere communities under a broad, long-term Hg gradient in field conditions, distinguishing the effects of Hg contamination and plant presence from bulk soil dynamics. Our findings reinforce the idea that plants actively recruit stress-adaptive microbial communities, which in turn mitigate Hg toxicity and foster long-term microbiome diversity. The identification of taxa such as *Septoglomus*, *Dioszegia*, and *Articulospora* as potential candidates for enhanced assisted-phytoremediation highlights key targets for future research into plant-microbe interactions that stabilize and detoxify Hg-contaminated soils.

## Acknowledgments

This work was supported by the Romanian National Authority for Scientific Research and Innovation, CNCS —UEFISCDI, project number PN-III-P2-2.1-PED-2019-5254, contract no. 390PED/2020. Emanuela D. Tiodar received an Erasmus+ Traineeship scholarship to conduct the bioinformatic analyses at the Biology Centre CAS, České Budějovice, Czech Republic. We would like to extend our warmest gratitude to PhD student Johannes Schweichhart for his kind support in sharing the blastn pipeline for taxonomy identification. This manuscript is a preprint and has not yet been peer-reviewed.

## Authors’ contributions

Conceptualization: EDT, DP, RA; Field sampling: ET, DP, ZRB, CT, AA; Formal analysis and investigation: all; Writing - original draft preparation: ET; Writing - review and editing: all; Funding acquisition: DP, ET; Resources: DP, RA, MB. Supervision: DP, RA, MB.

## Data availability

Sequence data generated in this study have been deposited in the European Nucleotide Archive (ENA) at EMBL-EBI under project accession numbers PRJEB88147 (data is private until journal acceptance). Supplementary data and figures are deposited in Zenodo: https://doi.org/10.5281/zenodo.15165625, and are currently under embargo. They will be made openly available upon journal acceptance.

## References

Abarenkov, K., Zirk, A., Piirmann, T., Pöhönen, R., Ivanov, F., Nilsson, R. H., & Kõljalg, U. (2024). Full UNITE+INSD dataset for Fungi. Version 21.04.2024. UNITE Community. 10.15156/BIO/2959330

Abdelaziz, A. M., El-Wakil, D. A., Hashem, A. H., Al-Askar, A. A., AbdElgawad, H., & Attia, M. S. (2023). Efficient Role of Endophytic *Aspergillus terreus* in Biocontrol of *Rhizoctonia solani* Causing Damping-off Disease of *Phaseolus vulgaris* and *Vicia faba*. Microorganisms, 11(6). 10.3390/microorganisms11061487

Abdu, N., Abdullahi, A. A., & Abdulkadir, A. (2017). Heavy metals and soil microbes. Environmental Chemistry Letters, 15(1), 65–84. 10.1007/s10311-016-0587-x

Agler, M. T., Ruhe, J., Kroll, S., Morhenn, C., Kim, S. T., Weigel, D., & Kemen, E. M. (2016). Microbial Hub Taxa Link Host and Abiotic Factors to Plant Microbiome Variation. PLoS Biology, 14(1). 10.1371/journal.pbio.1002352

Al-Huqail, A. A., & El-Bondkly, A. M. A. (2022). Improvement of *Zea mays* L. growth parameters under chromium and arsenic stress by the heavy metal-resistant *Streptomyces* sp. NRC21696. International Journal of Environmental Science and Technology, 19(3), 5301–5322. 10.1007/s13762-021-03532-7

Ali, A., Guo, D., Li, Y., Shaheen, S. M., Wahid, F., Antoniadis, V., Abdelrahman, H., Al-Solaimani, S. G., Li, R., Tsang, D. C. W., Rinklebe, J., & Zhang, Z. (2021). *Streptomyces pactum* addition to contaminated mining soils improved soil quality and enhanced metals phytoextraction by wheat in a green remediation trial. Chemosphere, 273, 129692. 10.1016/J.CHEMOSPHERE.2021.129692

Alori, E. T., Glick, B. R., & Babalola, O. O. (2017). Microbial Phosphorus Solubilization and Its Potential for Use in Sustainable Agriculture. Frontiers in Microbiology, 8. 10.3389/fmicb.2017.00971

Amaike, S., & Keller, N. P. (2011). Aspergillus flavus. Annual Review of Phytopathology, 49(Volume 49, 2011), 107–133. 10.1146/annurev-phyto-072910-095221

AMAP/UN, E. (2019). Technical Background Report for the Global Mercury Assessment 2018 (Arctic Monitoring and Assessment Programme, p. viii + 426 pp including E-Annexes). Chemicals and Health Branch, Geneva, Switzerland. https://www.amap.no/documents/doc/technical-background-report-for-the-global-mercury-assessment-2018/1815

An, D.-S., Im, W.-T., Yang, H.-C., & Lee, S.-T. (2006). *Shinella granuli* gen. Nov., sp. Nov., and proposal of the reclassification of *Zoogloea ramigera* ATCC 19623 as Shinella zoogloeoides sp. Nov. International Journal of Systematic and Evolutionary Microbiology, 56(Pt 2), 443–448. 10.1099/ijs.0.63942-0

Ankati, S., Srinivas, V., Pratyusha, S., & Gopalakrishnan, S. (2021). *Streptomyces* consortia-mediated plant defense against *Fusarium* wilt and plant growth-promotion in chickpea. Microbial Pathogenesis, 157, 104961. 10.1016/j.micpath.2021.104961

Anthony, M. A., Bender, S. F., & Heijden, M. G. A. van der. (2023). Enumerating soil biodiversity. Proceedings of the National Academy of Sciences of the United States of America, 120(33). 10.1073/pnas.2304663120

ATSDR. (2022). Toxicological Profile for Mercury (Draft for Public Comment). Atlanta, GA: Agency for Toxic Substances and Disease Registry (Issue April). https://www.atsdr.cdc.gov/toxprofiles/tp46.pdf

Bago, B., Pfeffer, P. E., & Shachar-Hill, Y. (2000). Carbon metabolism and transport in arbuscular mycorrhizas. Plant Physiology, 124(3). 10.1104/pp.124.3.949

Balázs, H. E., Schmid, C. A. O., Cruzeiro, C., Podar, D., Szatmari, P. M., Buegger, F., Hufnagel, G., Radl, V., & Schröder, P. (2021). Post-reclamation microbial diversity and functions in hexachlorocyclohexane (HCH) contaminated soil in relation to spontaneous HCH tolerant vegetation. Science of the Total Environment, 767, 144653. 10.1016/j.scitotenv.2020.144653

Balázs, H. E., Schmid, C. A. O., Feher, I., Podar, D., Szatmari, P. M., Marincaş, O., Balázs, Z. R., & Schröder, P. (2018). HCH phytoremediation potential of native plant species from a contaminated urban site in Turda, Romania. Journal of Environmental Management, 223(May), 286–296. 10.1016/j.jenvman.2018.06.018

Bertness, M. D., & Callaway, R. (1994). Positive interactions in communities. Trends in Ecology & Evolution, 9(5), 191–193. 10.1016/0169-5347(94)90088-4

Błaszkowski, J., Chwat, G., Góralska, A., Ryszka, P., & Orfanoudakis, M. (2014). *Septoglomus jasnowskae* and *Septoglomus turnauae*, two new species of arbuscular mycorrhizal fungi (Glomeromycota). Mycological Progress, 13(4), 985. 10.1007/s11557-014-0985-z

Bolchi, A., Ruotolo, R., Marchini, G., Vurro, E., Toppi, L. S. di, Kohler, A., Tisserant, E., Martin, F., & Ottonello, S. (2011). Genome-wide inventory of metal homeostasis-related gene products including a functional phytochelatin synthase in the hypogeous mycorrhizal fungus Tuber melanosporum. Fungal Genetics and Biology, 48(6), 573–584. 10.1016/J.FGB.2010.11.003

Bourdineaud, J. P., Durn, G., Režun, B., Manceau, A., & Hrenović, J. (2020). The chemical species of mercury accumulated by *Pseudomonas idrijaensis*, a bacterium from a rock of the Idrija mercury mine, Slovenia. Chemosphere, 248. 10.1016/j.chemosphere.2020.126002

Boyd, E. S., & Barkay, T. (2012). The mercury resistance operon: From an origin in a geothermal environment to an efficient detoxification machine. Frontiers in Microbiology, 3(OCT), 1–13. 10.3389/fmicb.2012.00349

Callahan, B. J., McMurdie, P. J., Rosen, M. J., Han, A. W., Johnson, A. J. A., & Holmes, S. P. (2016). DADA2: High-resolution sample inference from Illumina amplicon data. Nature Methods, 13(7), 581–583. 10.1038/nmeth.3869

Carrasco-Gil, S., Siebner, H., Leduc, D. L., Webb, S. M., Millán, R., Andrews, J. C., & Hernández, L. E. (2013). Mercury localization and speciation in plants grown hydroponically or in a natural environment. Environmental Science and Technology, 47(7), 3082–3090. 10.1021/es303310t

Chang, J., Si, G., Dong, J., Yang, Q., Shi, Y., Chen, Y., Zhou, K., & Chen, J. (2021). Transcriptomic analyses reveal the pathways associated with the volatilization and resistance of mercury(II) in the fungus *Lecythophora* sp. DC-F1. Science of the Total Environment, 752. 10.1016/j.scitotenv.2020.142172

Christakis, C. A., Barkay, T., & Boyd, E. S. (2021). Expanded Diversity and Phylogeny of *mer* Genes Broadens Mercury Resistance Paradigms and Reveals an Origin for MerA Among Thermophilic Archaea. Frontiers in Microbiology, 12(June), 1–20. 10.3389/fmicb.2021.682605

Crosbie, D. B., Mahmoudi, M., Radl, V., Brachmann, A., Schloter, M., Kemen, E., & Marín, M. (2022). Microbiome profiling reveals that *Pseudomonas antagonises* parasitic nodule colonisation of cheater rhizobia in *Lotus*. New Phytologist, 234(1), 242–255. 10.1111/nph.17988

Davis, N. M., Proctor Di. M., Holmes, S. P., Relman, D. A., & Callahan, B. J. (2018). Simple statistical identification and removal of contaminant sequences in marker-gene and metagenomics data. Microbiome, 6(1). 10.1186/s40168-018-0605-2

Durand, A., Maillard, F., Foulon, J., & Chalot, M. (2020, December). Interactions between Hg and soil microbes: Microbial diversity and mechanisms, with an emphasis on fungal processes. In Applied Microbiology and Biotechnology (Vol. 104, Issue 23, pp. 9855–9876). Springer Science and Business Media Deutschland GmbH. 10.1007/s00253-020-10795-6

Durand, A., Maillard, F., Foulon, J., Gweon, H. S., Valot, B., & Chalot, M. (2017). Environmental Metabarcoding Reveals Contrasting Belowground and Aboveground Fungal Communities from Poplar at a Hg Phytomanagement Site. Microbial Ecology, 74(4), 795–809. 10.1007/s00248-017-0984-0

Egidi, E., Delgado-Baquerizo, M., Plett, J. M., Wang, J., Eldridge, D. J., Bardgett, R. D., Maestre, F. T., & Singh, B. K. (2019). A few Ascomycota taxa dominate soil fungal communities worldwide. Nature Communications, 10(1), 2369. 10.1038/s41467-019-10373-z

Esbrí, J. M., Cacovean, H., & Higueras, P. (2018). Usage Proposal of a common urban decorative tree (*Salix alba* L.) to monitor the dispersion of gaseous mercury: A case study from Turda (Romania). Chemosphere, 193, 74–81. 10.1016/j.chemosphere.2017.11.007

European Comission. (2011). Soil: The hidden part of the climate cycle. Publications Office. doi:10.2779/30669

European Comission. (2021). EU missions – Soil deal for Europe – Concrete solutions for our greatest challenges. Publications Office of the European Union. doi10.2777/247887

European Commission. (2016). Soil threats in Europe. Publications Office.

European Commission. (2020). Caring for soil is caring for life – Ensure 75% of soils are healthy by 2030 for healthy food, people, nature and climate – Interim report of the mission board for soil health and food (Issue KI-02-20-463-EN-C). 10.2777/918775

European Commission. (2023). EU efforts for sustainable soil management: Unambitious standards and limited targeting. Special report 19, 2023. 52.

FAO. (2022, July). Soils for nutrition: State of the art. Rome. 10.4060/cc0900en

FAO, & ITPS. (2015). Status of the World’s Soil Resources (SWSR) – Technical Summary. https://www.fao.org/fileadmin/user_upload/newsroom/docs/FAO-world-soils-report-SUMMARY.pdf

Feng, T., Liu, Y., Huang, M., Chen, G., Tian, Q., Duan, C., & Chen, J. (2024). Reshaping the root endophytic microbiota in plants to combat mercury-induced stress. Science of the Total Environment, 945. 10.1016/j.scitotenv.2024.174019

Frey, B., & Rieder, S. R. (2013). Response of forest soil bacterial communities to mercury chloride application. Soil Biology and Biochemistry, 65, 329–337. 10.1016/j.soilbio.2013.06.001

Frossard, A., Donhauser, J., Mestrot, A., Gygax, S., Bååth, E., & Frey, B. (2018). Long- and short-term effects of mercury pollution on the soil microbiome. Soil Biology and Biochemistry, 120(August 2017), 191–199. 10.1016/j.soilbio.2018.01.028

Frossard, A., Hartmann, M., & Frey, B. (2017). Tolerance of the forest soil microbiome to increasing mercury concentrations. Soil Biology and Biochemistry, 105, 162–176. 10.1016/j.soilbio.2016.11.016

Ginting, R. C. B., Solihat, N., Hafsari, A. R., & Irawan. (2021). Potential bacteria capable of remediating mercury contaminated soils. IOP Conference Series: Earth and Environmental Science, 648(1). 10.1088/1755-1315/648/1/012136

González, D., Robas, M., Fernández, V., Bárcena, M., Probanza, A., & Jiménez, P. A. (2022). Comparative Metagenomic Study of Rhizospheric and Bulk Mercury-Contaminated Soils in the Mining District of Almadén. Frontiers in Microbiology, 13(March). 10.3389/fmicb.2022.797444

Google Maps, 2023, Turda, Romania, retrieved on 4.08.2023, https://www.google.com/maps/place/Turda,+Romania/@46.5613859,23.7615103,10231m/data=!3m1!1e3!4m15!1m8!3m7!1s0x47496620aa78a95b:0xc7f433dbd893f8a3!2sTurda,+Romania!3b1!8m2!3d46.564676!4d23.7971063!16zL20vMDNxMDV6!3m5!1s0x47496620aa78a95b:0xc7f433dbd893f8a3!8m2!3d46.564676!4d23.7971063!16zL20vMDNxMDV6?entry=ttu

Google Maps, 2023, Fostele fabric chimice Turda, Turda, Romania, retrieved on 4.08.2023, https://www.google.com/maps/place/Fostele+fabrici+Chimice+Turda/@46.558422,23.7792582,116m/data=!3m1!1e3!4m6!3m5!1s0x47496881f0000001:0x68ee3987b1a49c90!8m2!3d46.5572413!4d23.7816733!16s%2Fg%2F11kj8_tx7x?entry=ttu

Google Maps, 2023, Salina Turda, retrieved on 4.08.2023, https://www.google.com/maps/place/Salina+Turda/@46.5873125,23.7915421,261m/data=!3m1!1e3!4m6!3m5!1s0x474968b385bfc665:0xd47e573765edc10c!8m2!3d46.5877459!4d23.7872654!16s%2Fm%2F0ds9t5w?entry=ttu

Gorordo, M. F., Lucca, M. E., & Sangorrín, M. P. (2022). Biocontrol Efficacy of the Vishniacozyma Victoriae in Semi-Commercial Assays for the Control of Postharvest Fungal Diseases of Organic Pears. Current Microbiology, 79(9), 259. 10.1007/s00284-022-02934-1

Gramisci, B. R., Lutz, M. C., Lopes, C. A., & Sangorrín, M. P. (2018). Enhancing the efficacy of yeast biocontrol agents against postharvest pathogens through nutrient profiling and the use of other additives. Biological Control, 121, 151–158. 10.1016/j.biocontrol.2018.03.001

Gueldry, O., Lazard, M., Delort, F., Dauplais, M., Grigoras, I., Blanquet, S., & Plateau, P. (2003). Ycf1p-dependent Hg(II) detoxification in *Saccharomyces cerevisiae*. European Journal of Biochemistry, 270(11), 2486–2496. 10.1046/J.1432-1033.2003.03620.X

Guillén, A., Serrano-Tamay, F. J., Peris, J. B., & Arrillaga, I. (2020). *Glomus ibericum*, *Septoglomus mediterraneum*, and *Funneliformis pilosus*, three new species of arbuscular mycorrhizal fungi. Mycologia, 819–828. 10.1080/00275514.2020.1771992

Guo, X., Zhang, X., Qin, Y., Liu, Y. X., Zhang, J., Zhang, N., Wu, K., Qu, B., He, Z., Wang, X., Zhang, X., Hacquard, S., Fu, X., & Bai, Y. (2020). Host-Associated Quantitative Abundance Profiling Reveals the Microbial Load Variation of Root Microbiome. Plant Communications, 1(1), 1–17. 10.1016/j.xplc.2019.100003

Guo, Y., Deng, W., Mo, Q., Yu, Y., Zhang, Z., Wei, M., Tang, R., Lu, S., & Su, Y. (2025). Glutathione reductase plays a role in the metabolism of methylmercury degradation in *Rhodotorula mucilaginosa*. Microbiology Spectrum, 13(2). 10.1128/SPECTRUM.02395-24

Gweon, H. S., Oliver, A., Taylor, J., Booth, T., Gibbs, M., Read, D. S., Griffiths, R. I., & Schonrogge, K. (2015). PIPITS: an automated pipeline for analyses of fungal internal transcribed spacer sequences from the Illumina sequencing platform. 10.1111/2041-210X.12399

Halo, B. A., Al-Yahyai, R. A., & Al-Sadi, A. M. (2018). Aspergillus terreus inhibits growth and induces morphological abnormalities in *Pythium aphanidermatum* and suppresses pythium-induced damping-off of cucumber. Frontiers in Microbiology, 9(FEB). 10.3389/fmicb.2018.00095

Handa, Y., Nishide, H., Takeda, N., Suzuki, Y., Kawaguchi, M., & Saito, K. (2015). RNA-seq Transcriptional Profiling of an Arbuscular Mycorrhiza Provides Insights into Regulated and Coordinated Gene Expression in *Lotus japonicus* and *Rhizophagus irregularis*. Plant and Cell Physiology, 56(8), 1490–1511. 10.1093/pcp/pcv071

Harrison, M. J., Dewbre, G. R., & Liu, J. (2002). A phosphate transporter from *Medicago truncatula* involved in the acquisition of phosphate released by arbuscular mycorrhizal fungi. Plant Cell, 14(10), 2413–2429. 10.1105/tpc.004861

Hemkemeyer, M., Schwalb, S. A., Heinze, S., Joergensen, R. G., & Wichern, F. (2021). Functions of elements in soil microorganisms. Microbiological Research, 252, 126832. 10.1016/J.MICRES.2021.126832

Holtze, M. S., Nielsen, P., Ekelund, F., Rasmussen, L. D., & Johnsen, K. (2006). Mercury affects the distribution of culturable species of *Pseudomonas* in soil. Applied Soil Ecology, 31(3), 228–238. 10.1016/j.apsoil.2005.05.004

Hou, L., Hernández-Restrepo, M., Groenewald, J. Z., Cai, L., & Crous, P. W. (2020). Citizen science project reveals high diversity in didymellaceae (pleosporales, dothideomycetes). MycoKeys, 65, 49–99. 10.3897/mycokeys.65.47704

Huang, X., Men, P., Tang, S., & Lu, X. (2021). *Aspergillus terreus* as an industrial filamentous fungus for pharmaceutical biotechnology. Current Opinion in Biotechnology, 69, 273–280. 10.1016/J.COPBIO.2021.02.004

Huang, Z., Wu, D., Yang, S., Fu, W., Ma, D., Yao, Y., Lin, H., Yuan, J., Yang, Y., & Zhuang, Z. (2023). Regulation of Fungal Morphogenesis and Pathogenicity of *Aspergillus flavus* by Hexokinase AfHxk1 through Its Domain Hexokinase_2. Journal of Fungi, 9(11). 10.3390/jof9111077

Ihrmark, K., Bödeker, I. T. M., Cruz-Martinez, K., Friberg, H., Kubartova, A., Schenck, J., Strid, Y., Stenlid, J., Brandström-Durling, M., Clemmensen, K. E., & Lindahl, B. D. (2012). New primers to amplify the fungal ITS2 region— Evaluation by 454-sequencing of artificial and natural communities. FEMS Microbiology Ecology, 82(3), 666–677. 10.1111/j.1574-6941.2012.01437.x

Ikenaga, M., Kataoka, M., Yin, X., Murouchi, A., & Sakai, M. (2021). Characterization and Distribution of Agar-degrading *Steroidobacter agaridevorans* sp. Nov., Isolated from Rhizosphere Soils. Microbes and Environments, 36(1). 10.1264/jsme2.ME20136

Jarvis, B. D. W., Pankhurst, C. E., & Patel, J. J. (1982). *Rhizobiurn loti*, a New Species of Legume Root Nodule Bacteria. In International Journal Of Systematic Bacteriology, 32(3), pp. 378–380.

Ji, H., Zhang, Y., Bararunyeretse, P., & Li, H. (2018). Characterization of microbial communities of soils from gold mine tailings and identification of mercury-resistant strain. Ecotoxicology and Environmental Safety, 165, 182–193. 10.1016/J.ECOENV.2018.09.011

Jin, H.-Q., Liu, H.-B., Xie, Y.-Y., Zhang, Y.-G., Xu, Q.-Q., Mao, L.-J., Li, X.-J., Chen, J., Lin, F.-C., & Zhang, C.-L. (2018). Effect of the dark septate endophytic fungus *Acrocalymma vagum* on heavy metal content in tobacco leaves. Symbiosis, 74(2), 89–95. 10.1007/s13199-017-0485-4

Kakouridis, A., Hagen, J. A., Kan, M. P., Mambelli, S., Feldman, L. J., Herman, D. J., Weber, P. K., Pett-Ridge, J., & Firestone, M. K. (2022). Routes to roots: Direct evidence of water transport by arbuscular mycorrhizal fungi to host plants. New Phytologist, 236(1), 210–221. 10.1111/nph.18281

Kavčič, A., Mikuš, K., Debeljak, M., Elteren, J. T. van, Arčon, I., Kodre, A., Kump, P., Karydas, A. G., Migliori, A., Czyzycki, M., & Vogel-Mikuš, K. (2019). Localization, ligand environment, bioavailability and toxicity of mercury in *Boletus* spp. And *Scutiger pes-caprae* mushrooms. Ecotoxicology and Environmental Safety, 184, 109623. 10.1016/j.ecoenv.2019.109623

Kinoshita, H., Sohma, Y., Ohtake, F., Ishida, M., Kawai, Y., Kitazawa, H., Saito, T., & Kimura, K. (2013). Biosorption of heavy metals by lactic acid bacteria and identification of mercury binding protein. Research in Microbiology, 164(7), 701–709. 10.1016/J.RESMIC.2013.04.004

Klich, M. A. (2007, November). *Aspergillus flavus*: The major producer of aflatoxin. In Molecular Plant Pathology (Vol. 8, Issue 6, pp. 713–722). 10.1111/j.1364-3703.2007.00436.x

Knapp, D. G., Kovács, G. M., Zajta, E., Groenewald, J. Z., & Crous, P. W. (2015). Dark septate endophytic pleosporalean genera from semiarid areas. Persoonia: Molecular Phylogeny and Evolution of Fungi, 35(1), 87–100. 10.3767/003158515X687669

Knapp, D. G., Pintye, A., & Kovács, G. M. (2012). The dark side is not fastidious—Dark septate endophytic fungi of native and invasive plants of semiarid sandy areas. PLoS ONE, 7(2). 10.1371/journal.pone.0032570

Ko, W.-H., Yang, C.-H., Lin, M.-J., Chen, C.-Y., & Tsou, Y.-J. (2011). *Humicola phialophoroides* sp. Nov. From soil with potential for biological control of plant diseases. Botanical Studies, 52, 197–202.

Krämer, U. (2010). Metal hyperaccumulation in plants. Annual Review of Plant Biology, 61, 517–534. 10.1146/annurev-arplant-042809-112156

Lang, J., Hu, J., Ran, W., Xu, Y., & Shen, Q. (2012). Control of cotton *Verticillium* wilt and fungal diversity of rhizosphere soils by bio-organic fertilizer. Biology and Fertility of Soils, 48(2), 191–203. 10.1007/s00374-011-0617-6

Lauber, C. L., Hamady, M., Knight, R., & Fierer, N. (2009). Pyrosequencing-based assessment of soil pH as a predictor of soil bacterial community structure at the continental scale. Applied and Environmental Microbiology, 75(15), 5111–5120. 10.1128/AEM.00335-09

Li, D., Li, X., Tao, Y., Yan, Z., & Ao, Y. (2022). Deciphering the bacterial microbiome in response to long-term mercury contaminated soil. Ecotoxicology and Environmental Safety, 229, 113062. 10.1016/j.ecoenv.2021.113062

Li, X., He, X., Hou, L., Ren, Y., Wang, S., & Su, F. (2018). Dark septate endophytes isolated from a xerophyte plant promote the growth of *Ammopiptanthus mongolicus* under drought condition. Scientific Reports, 8(1), 7896. 10.1038/s41598-018-26183-0

Lima, A. I. G., Corticeiro, S. C., & Figueira, E. M. de A. P. (2006). Glutathione-mediated cadmium sequestration in *Rhizobium leguminosarum*. Enzyme and Microbial Technology, 39(4), 763–769. 10.1016/J.ENZMICTEC.2005.12.009

Lin, D. X., Wang, E. T., Tang, H., Han, T. X., He, Y. R., Guan, S. H., & Chen, W. X. (2008). *Shinella kummerowiae* sp. Nov., a symbiotic bacterium isolated from root nodules of the herbal legume *Kummerowia stipulacea*. International Journal of Systematic and Evolutionary Microbiology, 58(Pt 6), 1409–1413. 10.1099/ijs.0.65723-0

Lin, H., & Peddada, S. D. (2020). Analysis of compositions of microbiomes with bias correction. Nature Communications, 11(1). 10.1038/s41467-020-17041-7

Liu, Y. R., He, J. Z., Zhang, L. M., & Zheng, Y. M. (2012). Effects of long-term fertilization on the diversity of bacterial mercuric reductase gene in a Chinese upland soil. Journal of Basic Microbiology, 52(1), 35–42. 10.1002/jobm.201100211

Liu, Y. R., Wang, J. J., Zheng, Y. M., Zhang, L. M., & He, J. Z. (2014). Patterns of Bacterial Diversity Along a Long-Term Mercury-Contaminated Gradient in the Paddy Soils. Microbial Ecology, 68(3), 575–583. 10.1007/s00248-014-0430-5

Lo, K. H., Lu, C. W., Liu, F. G., Kao, C. M., & Chen, S. C. (2022). Draft genome sequence of *Pseudomonas* sp. A46 isolated from mercury-contaminated wastewater. Journal of Basic Microbiology, 62(10), 1193–1201. 10.1002/JOBM.202200106

Lorenzo-Gutiérrez, D., Gómez-Gil, L., Guarro, J., Roncero, M. I. G., Fernández-Bravo, A., Capilla, J., & López-Fernández, L. (2019). Role of the *Fusarium oxysporum* metallothionein Mt1 in resistance to metal toxicity and virulence. Metallomics, 11(7), 1230–1240. 10.1039/C9MT00081J

Lorite, M. J., Estrella, M. J., Escaray, F. J., Sannazzaro, A., Castro, I. M. V. E., Monza, J., Sanjuán, J., & León-Barrios, M. (2018, September). The Rhizobia-Lotus symbioses: Deeply specific and widely diverse. In Frontiers in Microbiology (Vol. 9, Issue SEP). Frontiers Media S.A. 10.3389/fmicb.2018.02055

Louis, B., Waikhom, S. D., Roy, P., Bhardwaj, P. K., Singh, M. W., Chandradev, S. K., & Talukdar, N. C. (2014). Invasion of *Solanum tuberosum* L. by *Aspergillus terreus*: A microscopic and proteomics insight on pathogenicity. BMC Research Notes, 7, 350. 10.5061/dryad.590j0

Luo, G., Shi, Z., Wang, H., & Wang, G. (2012). *Skermanella stibiiresistens* sp. Nov., a highly antimony-resistant bacterium isolated from coal-mining soil, and emended description of the genus *Skermanella*. International Journal of Systematic and Evolutionary Microbiology, 62(Pt 6), 1271–1276. 10.1099/ijs.0.033746-0

Luo, J., Gu, S., Guo, X., Liu, Y., Tao, Q., Zhao, H. P., Liang, Y., Banerjee, S., & Li, T. (2022). Core Microbiota in the Rhizosphere of Heavy Metal Accumulators and Its Contribution to Plant Performance. Environmental Science and Technology, 56(18), 12975–12987. 10.1021/acs.est.1c08832

Ma, M., Du, H., Sun, T., An, S., Yang, G., & Wang, D. (2019). Characteristics of archaea and bacteria in rice rhizosphere along a mercury gradient. Science of the Total Environment, 650, 1640–1651. 10.1016/j.scitotenv.2018.07.175

Ma, Y., Wang, J., Liu, Y., Wang, X., Zhang, B., Zhang, W., Chen, T., Liu, G., Xue, L., & Cui, X. (2023). *Nocardioides*: “Specialists” for Hard-to-Degrade Pollutants in the Environment. Molecules, 28(21), Article 21. 10.3390/molecules28217433

Maghear, I. (2013). BILANT DE MEDIU Nivel I, SC CONSECVENT SRL, Turda, Cluj County, Romania (pp. 1–19).

Manceau, A., Nagy, K. L., Glatzel, P., & Bourdineaud, J. P. (2021). Acute Toxicity of Divalent Mercury to Bacteria Explained by the Formation of Dicysteinate and Tetracysteinate Complexes Bound to Proteins in Escherichia coli and Bacillus subtilis. Environmental Science and Technology, 55(6), 3612–3623. 10.1021/ACS.EST.0C05202

Marčiulynas, A., Marčiulynienė, D., Lynikienė, J., Bakys, R., & Menkis, A. (2022). Fungal Communities in Leaves and Roots of Healthy-Looking and Diseased *Ulmus glabra*. Microorganisms, 10(11). 10.3390/microorganisms10112228

Mariano, C., Mello, I. S., Barros, B. M., da Silva, G. F., Terezo, A. J., & Soares, M. A. (2020). Mercury alters the rhizobacterial community in Brazilian wetlands and it can be bioremediated by the plant-bacteria association. Environmental Science and Pollution Research, 27(12), 13550–13564. 10.1007/s11356-020-07913-2

Marschner, H. (2012). Marschner’s Mineral Nutrition of Higher Plants. Edition No. 3 (Issue 3).

Maurer, D., Malique, F., Alfarraj, S., Albasher, G., Horn, M. A., Butterbach-Bahl, K., Dannenmann, M., & Rennenberg, H. (2021). Interactive regulation of root exudation and rhizosphere denitrification by plant metabolite content and soil properties. Plant and Soil, 467(1), 107–127. 10.1007/s11104-021-05069-7

Ministerial Order, No. 756/1997. (1997). Approving the Regulation Concerning the Assessment of Environmental Pollution. (In Romanian). Official Gazette Part. I, No. 303bis/06.11.1997.

Mohamed, A. H., El-Megeed, F. H. A., Hassanein, N. M., Youseif, S. H., Farag, P. F., Saleh, S. A., Abdel-Wahab, B. A., Alsuhaibani, A. M., Helmy, Y. A., & Abdel-Azeem, A. M. (2022). Native Rhizospheric and Endophytic Fungi as Sustainable Sources of Plant Growth Promoting Traits to Improve Wheat Growth under Low Nitrogen Input. Journal of Fungi, 8(2). 10.3390/jof8020094

Montiel-Rozas, M. M., Madejón, E., & Madejón, P. (2016). Effect of heavy metals and organic matter on root exudates (low molecular weight organic acids) of herbaceous species: An assessment in sand and soil conditions under different levels of contamination. Environmental Pollution, 216, 273–281. 10.1016/J.ENVPOL.2016.05.080

Muryani, E., Sajidan, Budiastuti, M. T. S., & Pranoto. (2023). Diversity and potential of herbaceous plants as mercury (Hg) hyperaccumulators in small-scale gold mining sites in Pancurendang, Banyumas, Indonesia. Biodiversitas, 24(6), 3364–3372. 10.13057/biodiv/d240632

Nannipieri, P., Ascher, J., Ceccherini, M. T., Landi, L., Pietramellara, G., & Renella, G. (2017). Microbial diversity and soil functions. European Journal of Soil Science, 68(1), 12–26.

Narmani, A., Teponno, R. B., Helaly, S. E., Arzanlou, M., & Stadler, M. (2019). Cytotoxic, anti-biofilm and antimicrobial polyketides from the plant associated fungus *Chaetosphaeronema achilleae*. Fitoterapia, 139, 104390. 10.1016/j.fitote.2019.104390

Natasha, Shahid, M., Khalid, S., Bibi, I., Bundschuh, J., Niazi, N. K., & Dumat, C. (2020). A critical review of mercury speciation, bioavailability, toxicity and detoxification in soil-plant environment: Ecotoxicology and health risk assessment. Science of the Total Environment, 711, 134749. 10.1016/j.scitotenv.2019.134749

Ndinga-Muniania, C., Mueller, R. C., Kuske, C. R., & Porras-Alfaro, A. (2021). Seasonal variation and potential roles of dark septate fungi in an arid grassland. Mycologia, 113(6), 1181–1198. 10.1080/00275514.2021.1965852

Nian, L., Xie, Y., Zhang, H., Wang, M., Yuan, B., Cheng, S., & Cao, C. (2023). *Vishniacozyma victoriae*: An endophytic antagonist yeast of kiwifruit with biocontrol effect to *Botrytis cinerea*. Food Chemistry, 411, 135442. 10.1016/J.FOODCHEM.2023.135442

Oksanen, J., Blanchet, F. G., Friendly, M., Kindt, R., Legendre, P., McGlinn, D., & Wagner, H. (2017). Vegan: Community ecology package. R package version 2.4–4.2. https://cran.r-project.org/package=vegan

Oladipo, O. G., Awotoye, Olusegun Olufemi, Olayinka, Akin, Ezeokoli, Obinna Tobechukwu, Maboeta, Mark Steve, & and Bezuidenhout, C. C. (2016). Heavy metal tolerance potential of Aspergillus strains isolated from mining sites. Bioremediation Journal, 20(4), 287–297. 10.1080/10889868.2016.1250722

Olanrewaju, O. S., & Babalola, O. O. (2019). *Streptomyces*: Implications and interactions in plant growth promotion. Applied Microbiology and Biotechnology, 103(3), 1179–1188. 10.1007/s00253-018-09577-y

Oliveira, A., Pampulha, M. E., Neto, M. M., & Almeida, A. C. (2010). Mercury tolerant diazotrophic bacteria in a long-term contaminated soil. Geoderma, 154(3–4), 359–363. 10.1016/j.geoderma.2009.11.008

Pang, F., Manoj,·, Solanki, K., & Wang, Z. (2022). *Streptomyces* can be an excellent plant growth manager. World Journal of Microbiology and Biotechnology, 38(3), 193. 10.1007/s11274-022-03380-8

Park, J., Song, W. Y., Ko, D., Eom, Y., Hansen, T. H., Schiller, M., Lee, T. G., Martinoia, E., & Lee, Y. (2012). The phytochelatin transporters AtABCC1 and AtABCC2 mediate tolerance to cadmium and mercury. Plant Journal, 69(2), 278–288. 10.1111/j.1365-313X.2011.04789.x

Parniske, M. (2008). Arbuscular mycorrhiza: The mother of plant root endosymbioses. Nature Reviews Microbiology, 6(10), 763–775. 10.1038/nrmicro1987

Pietro-Souza, W., Pereira, F. de C., Mello, I. S., Stachack, F. F. F., Terezo, A. J., Cunha, C. N. da, White, J. F., Li, H., & Soares, M. A. (2020). Mercury resistance and bioremediation mediated by endophytic fungi. Chemosphere, 240. 10.1016/j.chemosphere.2019.124874

Põlme, S., Abarenkov, K., Nilsson, R. H., Lindahl, B. D., Clemmensen, K. E., Kauserud, H., Nguyen, N., Kjøller, R., Bates, S. T., Baldrian, P., Frøslev, T. G., Adojaan, K., Vizzini, A., Suija, A., Pfister, D., Baral, H.-O., Järv, H., Madrid, H., Nordén, J., … Tedersoo, L. (2021). FungalTraits: A user-friendly traits database of fungi and fungus-like stramenopiles. Fungal Diversity. 10.1007/s13225-020-00466-2

Porras-Alfaro, A., Herrera, J., Natvig, D. O., Lipinski, K., & Sinsabaugh, R. L. (2011). Diversity and distribution of soil fungal communities in a semiarid grassland. Mycologia, 103(1), 10–21. 10.3852/09-297

Prăvălie, R., Borrelli, P., Panagos, P., Ballabio, C., Lugato, E., Chappell, A., Miguez-Macho, G., Maggi, F., Peng, J., Niculiță, M., Roșca, B., Patriche, C., Dumitrașcu, M., Bandoc, G., Nita, I.-A., & Birsan, M.-V. (2024). A unifying modelling of multiple land degradation pathways in Europe. Nature Communications, 15(1), 3862. 10.1038/s41467-024-48252-x

Prosenkov, A., Cagnon, C., Gallego, J. L. R., & Pelaez, A. I. (2023). The microbiome of a brownfield highly polluted with mercury and arsenic. Environmental Pollution, 323. 10.1016/j.envpol.2023.121305

Puglisi, I., Faedda, R., Sanzaro, V., Piero, A. R. L., Petrone, G., & Cacciola, S. O. (2012). Identification of differentially expressed genes in response to mercury I and II stress in *Trichoderma harzianum*. Gene, 506(2), 325–330. 10.1016/J.GENE.2012.06.091

Quiñones, M. A., Ruiz-Díez, B., Fajardo, S., López-Berdonces, M. A., Higueras, P. L., & Fernández-Pascual, M. (2013). *Lupinus albus* plants acquire mercury tolerance when inoculated with an Hg-resistant *Bradyrhizobium* strain. Plant Physiology and Biochemistry, 73, 168–175. 10.1016/j.plaphy.2013.09.015

R Core Team. (2024). *_*R: A Language and Environment for Statistical Computing_. R Foundation for Statistical Computing. https://www.R-project.org/

Raimondo, M. L., & Carlucci, A. (2018). Characterization and pathogenicity assessment of *Plectosphaerella* species associated with stunting disease on tomato and pepper crops in Italy. Plant Pathology, 67(3), 626–641. 10.1111/ppa.12766

Raspanti, E., Cacciola, S. O., Gotor, C., Romero, L. C., & García, I. (2009). Implications of cysteine metabolism in the heavy metal response in *Trichoderma harzianum* and in three *Fusarium* species. Chemosphere, 76(1), 48–54. 10.1016/J.CHEMOSPHERE.2009.02.030

Reeves, R. D., Baker, A. J. M., Jaffré, T., Erskine, P. D., Echevarria, G., & Ent, A. van der. (2018). A global database for plants that hyperaccumulate metal and metalloid trace elements. In New Phytologist (Vol. 218, Issue 2). 10.1111/nph.14907

Ruiz-Díez, B., Quiñones, M. A., Fajardo, S., López, M. A., Higueras, P., & Fernández-Pascual, M. (2012). Mercury-resistant rhizobial bacteria isolated from nodules of leguminous plants growing in high Hg-contaminated soils. Applied Microbiology and Biotechnology, 96(2), 543–554. 10.1007/s00253-011-3832-z

Saccá, M. L., Caracciolo, A. B., Lenola, M. D., & Grenni, P. (2017). Ecosystem Services Provided By Soil Microorganisms. In Soil Biological Communities and Ecosystem Resilience (pp. 9–24). Springer International Publishing. 10.1007/978-3-319-63336-7_2

Saeki, K., & Kouchi, H. (2000). The *Lotus* Symbiont, *Mesorhizobium loti*: Molecular Genetic Techniques and Application. Journal of Plant Research, 113(4), 457–465. 10.1007/PL00013956

Safari, F., Akramian, M., Salehi-Arjmand, H., & Khadivi, A. (2019). Physiological and molecular mechanisms underlying salicylic acid-mitigated mercury toxicity in lemon balm (*Melissa officinalis* L.). Ecotoxicology and Environmental Safety, 183(August), 109542. 10.1016/j.ecoenv.2019.109542

Sarao, S. K., Boothe, V., Das, B. K., Gonzalez-Hernandez, J. L., & Brözel, V. S. (2024). *Bradyrhizobium* and the soybean rhizosphere: Species level bacterial population dynamics in established soybean fields, rhizosphere and nodules. Plant and Soil, 1–16. 10.1007/S11104-024-06814-4

Schloter, M., Nannipieri, P., Sørensen, S. J., & Elsas, J. D. van. (2018). Microbial indicators for soil quality. Biology and Fertility of Soils, 54(1), 1–10. 10.1007/s00374-017-1248-3

Sepúlveda, X., Silva, D., Ceballos, R., Vero, S., López, M. D., & Vargas, M. (2022). Endophytic Yeasts for the Biocontrol of *Phlyctema vagabunda* in Apples. Horticulturae, 8(6), Article 6. 10.3390/horticulturae8060535

Sokal, R. R., & Rohlf, F. J. (2012). Biometry. The principles and practice of statistics in biological research. (fourth ed.). W.H. Freeman and Company. 10.2307/2412280

Subhash, Y., & Lee, S.-S. (2016a). *Shinella curvata* sp. Nov., isolated from hydrocarbon-contaminated desert sands. International Journal of Systematic and Evolutionary Microbiology, 66(10), 3929–3934. 10.1099/ijsem.0.001290

Subhash, Y., & Lee, S.-S. (2016b). *Skermanella rosea* sp. Nov., isolated from hydrocarbon-contaminated desert sands. International Journal of Systematic and Evolutionary Microbiology, 66(10), 3951–3956. 10.1099/ijsem.0.001293

Sun, L., Ma, Y., Wang, H., Huang, W., Wang, X., Han, L., Sun, W., Han, E., & Wang, B. (2018). Overexpression of PtABCC1 contributes to mercury tolerance and accumulation in *Arabidopsis* and poplar. Biochemical and Biophysical Research Communications, 497(4), 997–1002. 10.1016/J.BBRC.2018.02.133

Sun, X., Li, P., & Zheng, G. (2021). Cellular and subcellular distribution and factors influencing the accumulation of atmospheric Hg in *Tillandsia usneoides* leaves. Journal of Hazardous Materials, 414, 125529. 10.1016/J.JHAZMAT.2021.125529

Tedersoo, L., Hansen, K., Perry, B. A., & Kjøller, R. (2006). Molecular and morphological diversity of pezizalean ectomycorrhiza. New Phytologist, 170, 581–596. 10.1111/j.1469-8137.2006.01678.x

Tedersoo, L., May, T. W., Smith, M. E., Tedersoo, L., May, T. W., & Smith, M. E. (2009). Ectomycorrhizal lifestyle in fungi: Global diversity, distribution, and evolution of phylogenetic lineages. Mycorrhiza 2009 20:4, 20(4), 217–263. 10.1007/S00572-009-0274-X

Tiodar, E. D., Chiriac, C. M., Pošćić, F., Văcar, C. L., Balázs, Z. R., Coman, C., Weindorf, D. C., Banciu, M., Krämer, U., & Podar, D. (2024). Plant colonizers of a mercury contaminated site: Trace metals and associated rhizosphere bacteria. Plant and Soil. 10.1007/s11104-024-06552-7

Tiodar, E. D., Văcar, C. L., & Podar, D. (2021). Phytoremediation and microorganisms-assisted phytoremediation of mercury-contaminated soils: Challenges and perspectives. International Journal of Environmental Research and Public Health, 18(5), 1–38. 10.3390/ijerph18052435

Tiodar, E. D., Văcar, C. L., Grimm, M. C., Ganea, I. V., Balázs, Z. R., Abrudan, A. M., Timár, C., Tanțău, I., Banciu, M., Angel, R., Podar, D. (2025). Supplementary Data for Impact of Long-Term Mercury Contamination on the Rhizosphere Microbiota of Lotus tenuis: A Pathway to Resilience via Interkingdom Facilitation [Data set]. Zenodo. 10.5281/zenodo.15165625

Tomalka, J., Hunecke, C., Murken, L., Heckmann, T., Cronauer, C. C., Becker, R., Collignon, Q., Collins-Sowah, P. A., Crawford, M., Gloy, N., Hampf, A., Lotze-Campen, H., Malevolti, G., Maskell, G. M., Müller, C., Popp, A., Vodounhessi, M., Gornott, C., & Rockström, J. (2024). Stepping back from the precipice: Transforming land management to stay within planetary boundaries. Potsdam Institute for Climate Impact Research. https://publications.pik-potsdam.de/pubman/faces/ViewItemOverviewPage.jsp?itemId=item_30631

Tsiknia, M., Tsikou, D., Papadopoulou, K. K., & Ehaliotis, C. (2021). Multi-species relationships in legume roots: From pairwise legume-symbiont interactions to the plant – microbiome – soil continuum. FEMS Microbiology Ecology, 97(2), fiaa222. 10.1093/femsec/fiaa222

Uqab, B., Nazir, R., Ganai, B. A., & Rahi, P. (2024). Mercury-tolerant metalophiles: A bio tool for remediation of mercury (Hg) affected Environs. Process Safety and Environmental Protection, 191, 2074–2081. 10.1016/J.PSEP.2024.09.072

Urík, M., Hlodák, M., Mikušová, P., & Matúš, P. (2014). Potential of microscopic fungi isolated from mercury contaminated soils to accumulate and volatilize mercury(II). Water, Air, and Soil Pollution, 225(12). 10.1007/s11270-014-2219-z

US EPA, O. (2007). U.S. EPA Method 3051A: Microwave Assisted Acid Digestion of Sediments, Sludges, and Oils [Data and Tools]. https://www.epa.gov/esam/us-epa-method-3051a-microwave-assisted-acid-digestion-sediments-sludges-and-oils

Văcar, C. L., Covaci, E., Chakraborty, S., Li, B., Weindorf, D. C., Frențiu, T., Pârvu, M., & Podar, D. (2021). Heavy Metal-Resistant Filamentous Fungi as Potential Mercury Bioremediators. Journal of Fungi 2021, Vol. 7, Page 386, 7(5), 386. 10.3390/JOF7050386

Velázquez, E., García-Fraile, P., Ramírez-Bahena, M.-H., Rivas, R., & Martínez-Molina, E. (2017). Current Status of the Taxonomy of Bacteria Able to Establish Nitrogen-Fixing Legume Symbiosis. In A. Zaidi, M. S. Khan, & J. Musarrat (Eds.), Microbes for Legume Improvement (pp. 1–43). Springer International Publishing. 10.1007/978-3-319-59174-2_1

Wang, L., Zhang, K., Shareen, Yin, Z., Yu, L., Qiu, X., Zhou, S., Chen, R., & Wang, Q. (2024). Exploring the potential of *Aspergillus terreus* 2021, WLL-ISO: A dual-functional agent for heavy metal removal and self-aggregation in wastewater treatment. Separation and Purification Technology, 350, 127964. 10.1016/j.seppur.2024.127964

Wang, Y., Wang, L., Suo, M., Qiu, Z., Wu, H., Zhao, M., & Yang, H. (2022). Regulating Root Fungal Community Using *Mortierella alpina* for *Fusarium oxysporum* Resistance in *Panax ginseng*. Frontiers in Microbiology, 13. 10.3389/fmicb.2022.850917

Wani, Z. A., Kumar, A., Sultan, P., Bindu, K., Riyaz-Ul-Hassan, S., & Ashraf, N. (2017). *Mortierella alpina* CS10E4, an oleaginous fungal endophyte of *Crocus sativus* L. enhances apocarotenoid biosynthesis and stress tolerance in the host plant. Scientific Reports, 7(1), 8598. 10.1038/s41598-017-08974-z

Water Frame Directive, EP, CONSIL, 348 OJ L (2008). http://data.europa.eu/eli/dir/2008/105/oj/eng

Wei, H., He, X., Riccardo, B., Yang, Y., & Yuan, Z. (2021). *Stagonosporopsis rhizophilae* sp. Nov. (Didymellaceae, Pleosporales), a new rhizospheric soil fungus associated with *Populus deltoides* Marsh. Phytotaxa, 491(1), Article 1. 10.11646/phytotaxa.491.1.3

Wu, C., Tang, D., Dai, J., Tang, X., Bao, Y., Ning, J., Zhen, Q., Song, H., Leger, R. J. S., & Fang, W. (2022). Bioremediation of mercury-polluted soil and water by the plant symbiotic fungus *Metarhizium robertsii*. Proceedings of the National Academy of Sciences of the United States of America, 119(47), e2214513119. 10.1073/PNAS.2214513119

Xavier, J. C., Costa, P. E. S., Hissa, D. C., Melo, V. M. M., Falcão, R. M., Balbino, V. Q., Mendonça, L. A. R., Lima, M. G. S., Coutinho, H. D. M., & Verde, L. C. L. (2019). Evaluation of the microbial diversity and heavy metal resistance genes of a microbial community on contaminated environment. Applied Geochemistry, 105, 1–6. 10.1016/j.apgeochem.2019.04.012

Zappelini, C., Karimi, B., Foulon, J., Lacercat-Didier, L., Maillard, F., Valot, B., Blaudez, D., Cazaux, D., Gilbert, D., Yergeau, E., Greer, C., & Chalot, M. (2015). Diversity and complexity of microbial communities from a chlor-alkali tailings dump. Soil Biology and Biochemistry, 90, 101–110. 10.1016/J.SOILBIO.2015.08.008

Zhang, Y., Zhang, Y., Liu, M., Shi, X., & Zhao, Z. (2008). Dark septate endophyte (DSE) fungi isolated from metal polluted soils: Their taxonomic position, tolerance, and accumulation of heavy metals In Vitro. The Journal of Microbiology, 46(6), 624– 632. 10.1007/s12275-008-0163-6

Zheng, X., Cao, H., Liu, B., Zhang, M., Zhang, C., Chen, P., & Yang, B. (2022). Effects of Mercury Contamination on Microbial Diversity of Different Kinds of Soil. Microorganisms, 10(5). 10.3390/microorganisms10050977

Zhu, W., Huang, J., Li, M., Li, X., & Wang, G. (2014). Genomic analysis of *Skermanella stibiiresistens* type strain SB22T. Standards in Genomic Sciences, 9(3), Article 3. 10.4056/sigs.5751047

